# Inferring cascade drivers of VEXAS syndrome by a causal machine learning tool CauNagi

**DOI:** 10.64898/2026.09.23.753348

**Authors:** Dengyi Bao, Han Yao, Jingjing Liu, Ge Dong, Shuqian Xu, Zhigang Cai

**Affiliations:** Department of Bioinformatics, State Key Laboratory of Experimental Hematology, Tianjin Key Laboratory of Inflammatory Biology, School of Basic Medical Science, Tianjin Medical University, Tianjin, China; Department of Pharmacology, The Province and Ministry Co-sponsored Collaborative Innovation Center for Medical Epigenetics, School of Basic Medical Science, Tianjin Medical University, Tianjin, China; Shandong Provincial Clinical Research Center for Hematological Diseases, Shandong Key Laboratory of Hematological Diseases and Immune Microenvironment, Jinan, China; Department of Hematology, Qilu Hospital of Shandong University, Jinan, Shandong, People’s Republic of China

**Keywords:** Causal machine learning, CauNagi, VEXAS syndrome, Hematopoietic hierarchy, Global candidate regulators, Cascade candidate regulators, Single-cell transcriptomics

## Abstract

VEXAS syndrome is an adult-onset severe autoinflammatory disease caused by somatic mutations in *UBA1*, yet the cascade mechanisms linking primitive hematopoietic abnormalities to mature myeloid dysfunctions remain largely unknown. Identifying master regulators of a progressive disease, a black-box process, from complex transcriptomic data also remains challenging. To address this challenge, we developed CauNagi, a computational framework for prioritizing cascade candidate regulators (CCRs). CauNagi integrates a causal representation learning module derived from CausCell with an iterative deep learning backbone adapted from UNAGI; in addition, CauNagi extends these two components with a unique downstream module for CCRs analysis designed to characterize regulatory propagation across hierarchical cellular states. Mechanistically, CauNagi iteratively integrates causal disentangled representation learning with (1) disease-stage cell-state trajectory reconstruction and (2) dynamic regulatory analysis. Benchmarking on single-cell transcriptomic datasets showed that CauNagi preserved cell-type structure in idiopathic pulmonary fibrosis (IPF) and enriched known acute myeloid leukemia (AML)- associated genes among its top-ranked global regulators. When applied to VEXAS syndrome, CauNagi readily revealed inflammatory responses, endoplasmic reticulum stress, and myeloid bias, consistent with the disease features. Furthermore, the CCRs analysis module of CauNagi assisted us in identifying 36 causal drivers, with *SPI1, NFKB1, STAT3*, and *FOS* prioritized as high-confidence regulatory hubs linking aberrant myeloid differentiation and inflammatory programs. These findings were further supported by an independent single-cell transcriptomic dataset from a murine VEXAS model. Overall, CauNagi provides a computationally efficient and systematic framework for identifying candidate causal regulators. Beyond hematopoietic diseases, CauNagi may also be applicable to other progressive disorders for which multistage single-cell transcriptomic datasets are available. CauNagi is available at https://github.com/steamed-stuffedbun/CauNagi.

**Highlights:**

1. CauNagi identifies disease-progression regulators from multistage single-cell transcriptomes and prioritizes cascade candidate regulators (CCRs) by comparing them along hematopoietic myeloid hierarchy;
2. Benchmark studies demonstrate strong performance of CauNagi in both latent representation learning and identification of global candidate regulators (GCRs)
3. To apply CauNagi for understanding VEXAS syndrome, we constructed a single-cell transcriptomic atlas of VEXAS syndrome, comprising 290,441 cells from 38 samples;
4. Cascade-based prioritization identified 36 core CCRs across myeloid differentiation trajectories, with supports from the VEXAS-like murine models developed in house;

## Introduction

The Vacuoles, E1 enzyme, X-linked, Autoinflammatory, Somatic (VEXAS) syndrome is a recently identified adult-onset autoinflammatory and hematologic disorder caused by somatic mutations in *UBA1*, an X-linked gene encoding ubiquitin-like modifier-activating enzyme 1.^1^ The disease predominantly affects older men and is characterized by persistent systemic inflammation, cytopenia, bone marrow dysplasia, and multiorgan involvement.^2^ Although the initiating genetic lesion is well established, the regulatory processes linking *UBA1* dysfunction to the heterogeneous inflammatory and hematopoietic manifestations of VEXAS remain unresolved.^3–4^ Defective *UBA1* activity, primarily driven by the loss of the cytoplasmic isoform UBA1b, disrupts cytoplasmic ubiquitination and proteostasis, producing abnormalities that are detectable in hematopoietic stem and progenitor cells (HSPCs) and become increasingly associated with myeloid bias, inflammatory activation, and cell death programs in downstream myeloid populations.^5–8^ The previous studies support a model in which the molecular consequences of *UBA1* dysfunction extend across successive stages of myeloid differentiation.^7–10^ However, the master regulators that connect early hematopoietic abnormalities to late and terminal inflammatory phenotypes have not been systematically identified.

Most recently multi-cohort single-cell studies of VEXAS provide stage-resolved observations and insights across hematopoiesis stem cell (HSCs), granulocyte-monocyte progenitors (GMPs), monocytes, and neutrophils,^4–5,7,9^ but they also introduce a distinct computational challenge: distinguishing main regulators that profoundly initiate, sustain, or amplify disease-associated programs from those that minorly reflect downstream inflammation. In addition, analysis from differentially expressed genes (DEGs) and pathway enrichment (PE) reveals molecular alterations but they are too simple and insufficient to explain how regulatory signals persist across cell states or identify key drivers among huge numbers of DEGs. ^11–12^ Trajectory-based methods reconstruct continuous transitions and identify dynamically expressed genes, yet expression dynamics alone do not determine which factors organize those transitions.^13,14^ Gene regulatory networks (GRNs) with driver gene hubs provide alternative and complementary information about transcriptional control, but many approaches depend on fixed prior networks or additional omic layers and may characterize regulatory relationships only within a single static state (a snapshot of an omic dataset).^15–18^ Moreover, comparisons across tissues, donors, cohorts, and sequencing protocols require disease-associated variation to be separated from cell identity and residual heterogeneity before regulatory evidence can be integrated.^19^ Therefore, it is necessary to develop an analytical framework for identifying key regulators across different disease stages. Such a framework should be able to jointly reconstruct cellular states, infer dynamic regulatory relationships, and quantify the cross-stage persistence of regulatory support within an ordered developmental hierarchy, such as the biological process of hematopoiesis.

To address these challenges, we developed a computational framework termed CauNagi. CauNagi builds on the causal representation strategy of CausCell and the iterative disease-progression analysis framework of UNAGI.^20,21^ At its core module, CauNagi iteratively couples causally disentangled representation learning with cross-stage cell-state matching and dynamic regulatory inference, and further incorporates a downstream module for prioritizing and ranking CCRs. The central premise of this module is that the potential cascade-regulatory role of a regulator is determined not only by its importance within an individual cell state, but also by whether and how its regulatory influence persists, propagates downstream, or changes dynamically across successive cell states. By explicitly modeling these cross-stage regulatory patterns, CauNagi enables candidate regulators to be evaluated across disease progression and thereby prioritizes key regulatory factors that may connect early molecular abnormalities to downstream pathological phenotypes. Such cross-stage regulatory continuity and propagation are particularly relevant to hematopoietic disorders and may be especially important for understanding the pathogenesis of VEXAS syndrome.

## Results

### Overview of the CauNagi framework

CauNagi was designed to identify candidate regulators that connect different disease states rather than to screen genes solely based on a single static comparison (snapshot). CauNagi takes two major inputs: 1) single-cell RNA-seq expression matrices from multiple disease stages; and 2) a causal directed acyclic graph (cDAG) constructed based on user-defined concepts and their corresponding labels (**Figure 1A**). The observable concepts include disease stage, cell identity, and other study covariates. Meanwhile, an additional unobserved concept is introduced to capture residual variation that cannot be explained by the available labels. This design enables the latent representation to preserve biologically interpretable information while preventing unexplained heterogeneity from being incorrectly assigned to predefined concepts.

**Figure 1:**
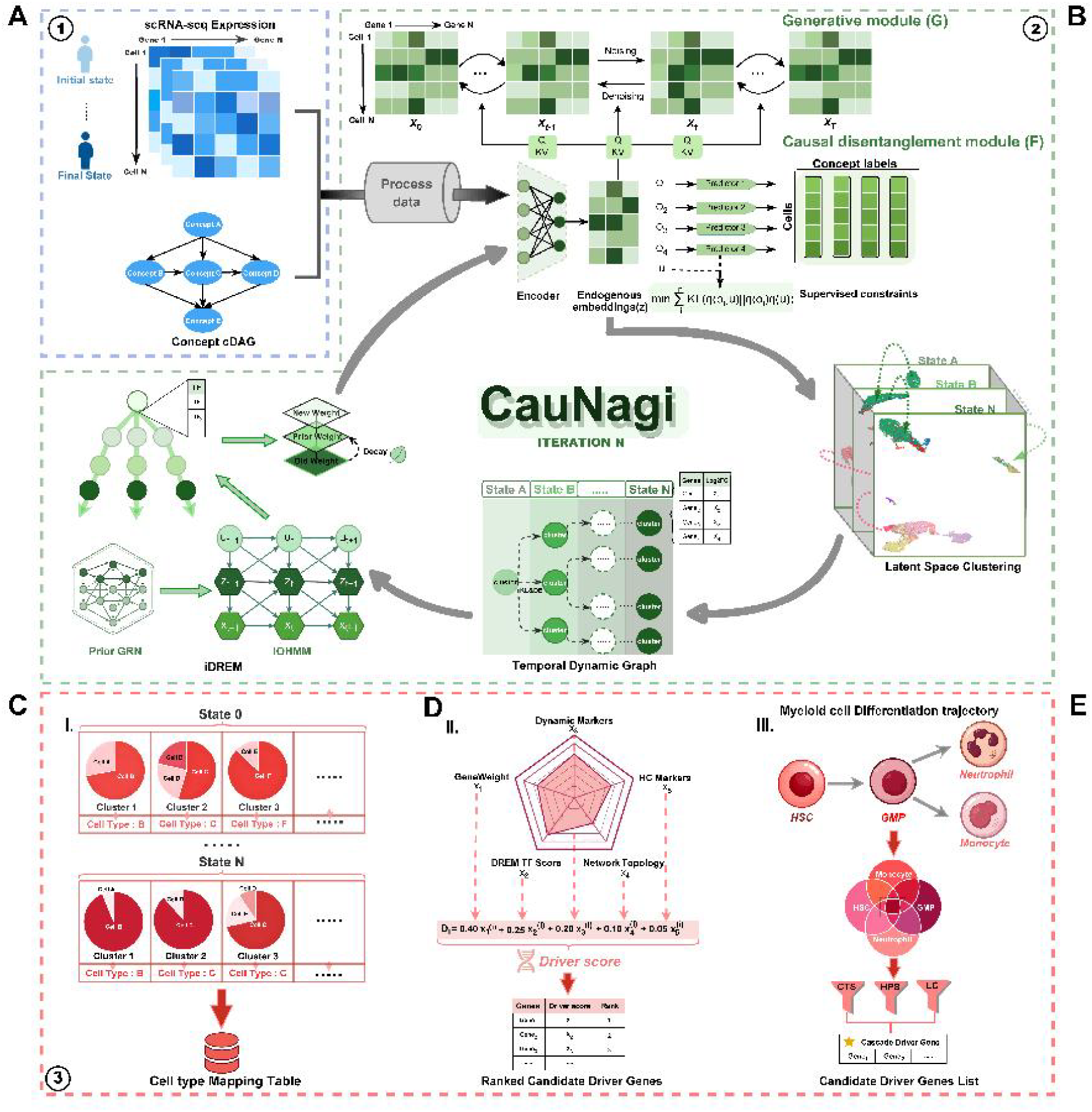
Architecture and downstream analytical modules of CauNagi. **(A)** CauNagi takes disease-stage single-cell RNA sequencing (scRNA-seq) expression matrices and a concept-level causal directed acyclic graph (cDAG) as inputs. The cDAG comprises observable concepts, such as disease stage and cell identity, together with an unexplained concept that captures residual heterogeneity. **(B)** CauNagi consists of two core modules: representation learning and dynamic feedback. The representation-learning module, inspired by CausCell, learns robust expression representations through concept-conditioned diffusion and causal disentanglement. Exogenous latent variables are mapped through a fixed concept-level causal graph to endogenous concept representations, separating known biological concepts from residual variation and providing a latent space for within-stage cell-state clustering. The dynamicfeedback module, inspired by UNAGI, connects cell states across adjacent stages into a temporally ordered dynamic graph, integrates iDREM with prior regulatory networks to extract dynamic TF–target evidence, and converts this evidence into gene weights that are fed back into the next training round, thereby enabling iterative coordination between latent representation learning and regulatory inference. **(C)** Generation of outputs cell-type information. Cell type mapping assigns latent clusters to cell identities across disease states and produces a cell type mapping table. **(D)** Gene-scoring module. Iterative gene weights, dynamic markers, iDREM transcriptionfactor evidence, network topology, and hierarchical-cluster markers are integrated into a driver score and ranked candidate list. **(E)** A downstream module of CauNagi, for CCRs analysis. Cell-type-specific candidates are evaluated along the HSPC-GMP-monocyte/neutrophil differentiation axis to derive CTS, HPS, and LC and to define candidate cascade driver genes.

The central part of CauNagi couples a concept-conditioned generative diffusion module to a causal disentanglement encoder (**Figure 1B**). The diffusion component learns a noise-robust representation of the expression distribution, whereas supervised concept predictors encourage separate latent subspaces to encode the corresponding biological labels. An adversarial constraint reduces leakage of known concepts into the unexplained component. Cells are subsequently clustered within each disease stage in the latent space, and state specific clusters are linked across adjacent stages by combining features in their latent distributions with distances between cluster marker profiles. The resulting temporal dynamic graph provides stage-aware and ordered expression trajectories for iDREM,^22^ which in turn integrates dynamic expression with a prior gene regulatory network (GRN) to identify effective transcription factors (TF) and candidate targets.

Secondly, during the machine learning process, regulatory evidence recovered in each iteration was fed back to the gene weight layer of the model. Genes supported by stage-dependent expression, transcription factor evidence, or prior TF target relationships will receive higher reconstruction weights in the next round, creating a closed loop between representation learning (RL) and dynamic regulatory inference. After convergence, CauNagi integrated iterative gene weights, iDREM evidence, dynamic markers, network topology, and hierarchical cluster markers into a unified score of filtered drivers. The final outputs included a cell type mapping table, ranked candidate regulators, and lineage-aware CCRs along the HSPC-GMP-monocyte/neutrophil differentiation axis (**Figure 1C-E**). Thus, CauNagi links cell state representation, temporal structure, and regulatory prioritization within one iterative analysis.

Of note, the original “head” module of UNAGI, the GAN-VAE, was replaced with Causal.VAE in CauNagi. We found that Causal.VAE could be seamlessly integrated with the remaining components of the UNAGI framework without disrupting its iterative learning architecture. Furthermore, the resulting CauNagi framework outperformed the original UNAGI framework in our benchmarking analyses, as shown below.

### Benchmark Evaluation of CauNagi on Single-cell Datasets

The performance of CauNagi depends on both informative cell representations and reliable regulator prioritization. These two underlying capabilities were therefore evaluated separately in the following studies.

We first assessed representation learning of CauNagi using the IPF single-cell dataset GSE286182, in which established cell type annotations provided a reference for evaluating whether the learned latent space preserved biologically meaningful cellular structure. CauNagi was compared with UNAGI,^21^ scGNN,^23^ scGGAN,^24^ GraphSCC,^25^ scGEN,^26^ scVI,^27^ scGPT,^28^ and Geneformer.^29^ These methods respectively represent graph-based, generative, probabilistic, and pretrained single-cell representation learning strategies. All methods were evaluated using matched inputs across ten independent runs and assessed with the scIB benchmark.^30^

CauNagi achieved the highest mean adjusted rand index (ARI) and normalized mutual information (NMI) among the compared methods, indicating strong agreement between the latent space organization and the reference cell type annotations (**Figure 2A**). It also maintained favorable performance across complementary measures of label preservation, neighborhood structure, and biological conservation. The aggregate metric profile showed that this performance was not driven by a single evaluation criterion but reflected balanced preservation of both local and global cellular structure (**Figure 2B**). Although CauNagi was not uniformly superior for every individual silhouette or isolated-label metric, its repeated run distributions remained stable, supporting the suitability of the learned representation for downstream state matching and trajectory construction.

**Figure 2:**
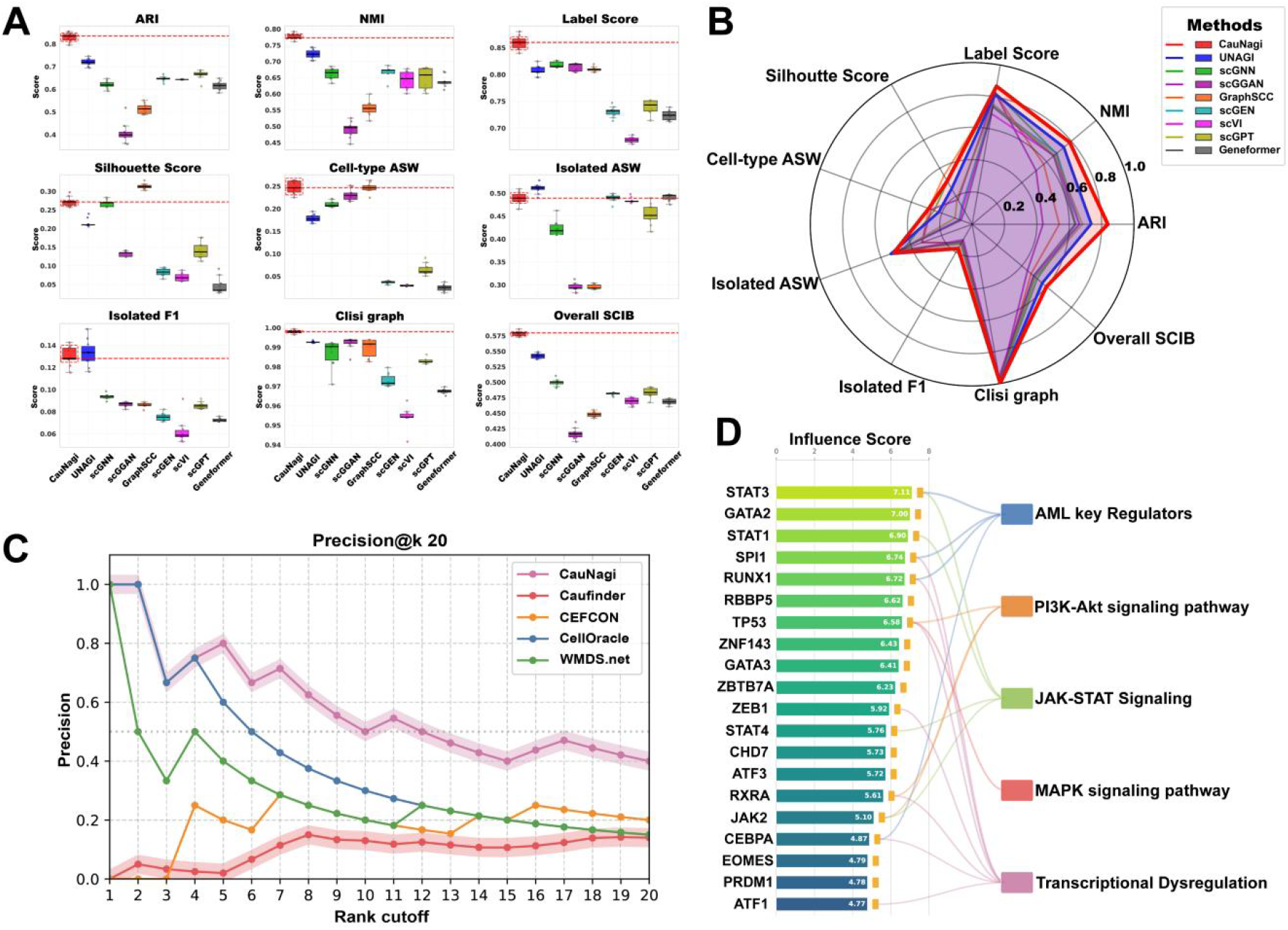
Benchmarking CauNagi in performances for latent space representation and driver-gene prioritization. **(A)** Boxplots of ARI, NMI, label score, silhouette score, cell type ASW, isolated-label ASW, isolated-label F1, graph cLISI, and overall scIB bio-conservation score across ten independent runs of CauNagi, UNAGI, scGNN, scGGAN, GraphSCC, scGEN, scVI, scGPT, and Geneformer on the IPF dataset GSE286182. Red dashed lines indicate CauNagi means. **(B)** Radar plot comparing mean benchmark profiles across the nine scIB metrics. CauNagi is highlighted in red. **(C)** Mean Precision@k for k = 1–20 across ten runs of CauNagi, CauFinder, CEFCON, CellOracle, and WMDS.net in the AML dataset GSE116256. Shaded bands indicate between-run dispersion. **(D)** Top 20 CauNagi AML candidates ranked by driver score and connected to the curated AML key-regulator set, PI3K-Akt signaling, JAK-STAT signaling, MAPK signaling, and transcriptional dysregulation.

We next investigated whether this performance could be attributed to the principal components of CauNagi and performed an ablation test. Removing causal disentanglement reduced ARI and NMI to near-zero levels and substantially impaired the remaining structure preservation metrics. By comparison, disabling iterative gene weight feedback produced a smaller but consistent deterioration in cluster separation and overall embedding quality (**Supplementary Fig. S1**). These results indicate that causal disentanglement provides the principal signal for separating biologically defined cell states, whereas iterative feedback further refines the representation by emphasizing genes supported by dynamic regulatory evidence.

We then evaluated the second core capability of CauNagi—candidate regulator prioritization—using the AML single-cell dataset GSE116256.^31^ An independent 36-gene set of the AML reference was assembled from DepMap functional dependency evidence,^32^ DisGeNET disease associations,^33^ published experimental studies, and human-expert review (**Supplementary Table 2**). CauNagi was compared with CellOracle,^15^ CEFCON,^17^ and WMDS.net,^18^ CauFinder ^34^ across ten independent runs. Because these methods differ in their model structures, prior network requirements, and definitions of regulatory importance, the comparison was restricted to their common output: ranked candidate gene lists. The benchmark therefore evaluated the recovery of independently supported AML regulators rather than the equivalence of the inferred networks or their causal interpretations.

CauNagi maintained high precision at the smallest ranking cutoffs and achieved the leading mean Precision@k from approximately the top five cutoff onward. At k = 20, it recovered eight reference regulators among its first 20 predictions, corresponding to a mean Precision@20 of 0.40 (**Figure 2C**). Candidate set comparison showed that the five methods generated partially overlapping but substantially method specific predictions, indicating that comparable list sizes did not necessarily result in equivalent recovery of the AML reference genes (**Supplementary Fig. S2**). Across both the top20 predictions and the complete reported candidate sets, CauNagi achieved the most favorable overall balance among precision, recall, and F1 score (**Supplementary Fig. S3**).

The leading CauNagi predictions included established hematopoietic and leukemia-associated regulators such as *STAT3, GATA2, SPI1, RUNX1*, and *CEBPA*,^35–38^ with the complete candidate list provided in **Supplementary Table 3**. Functional mapping further linked the top ranked candidates to AML-associated regulatory processes, including JAK– STAT, PI3K–Akt, and MAPK signaling, as well as transcriptional dysregulation (**Figure 2D**).^38–40^ The agreement among rank-based recovery, aggregate predictive performance, and pathway level interpretation supports the ability of CauNagi to prioritize biologically relevant disease-associated regulators.

Collectively, the benchmark studies with IPF and AML datasets provide direct evidence that CauNagi can learn biologically structured latent representations and generate reliable candidate regulator pools. These capabilities establish an empirical foundation for dissecting drivers in VEXAS syndrome which is complexed with dynamic hematopoietic hierarchies and branches.

### A multi-source single-cell atlas resolves VEXAS-associated cell states across myeloid differentiation

Most recently multi-cohort single-cell studies of VEXAS provide stage-resolved observations and insights across HSPCs, granulocyte-monocyte progenitors (GMPs), monocytes, and neutrophils.^4–5,7^ We integrated four publicly available VEXAS single-cell datasets—GSE272816,^4^ GSE196052,^5^ GSE216548,^7^ and GSE249131,^9^ with three in-house healthy bone marrow samples (**Figure 3A**). ^41^ The initial collection comprised 40 samples from 26 patients with VEXAS and 14 healthy controls, spanning bone marrow mononuclear cells (BMMC), CD34-enriched bone marrow cells, peripheral blood mononuclear cells (PBMC), and whole blood. Following stringent quality control, Samples marked as bm2.BM and PT9 were excluded because of abnormally high hemoglobin transcript fractions. Dataset specific filtering and doublet removal ultimately retained 290,441 high quality cells from 38 samples. ^42^ Harmony correction reduced sample associated separation in principal component space, and clustering of the corrected representation yielded 40 initial clusters for subsequent annotation (**Supplementary Fig. S4**).^43^

**Figure 3:**
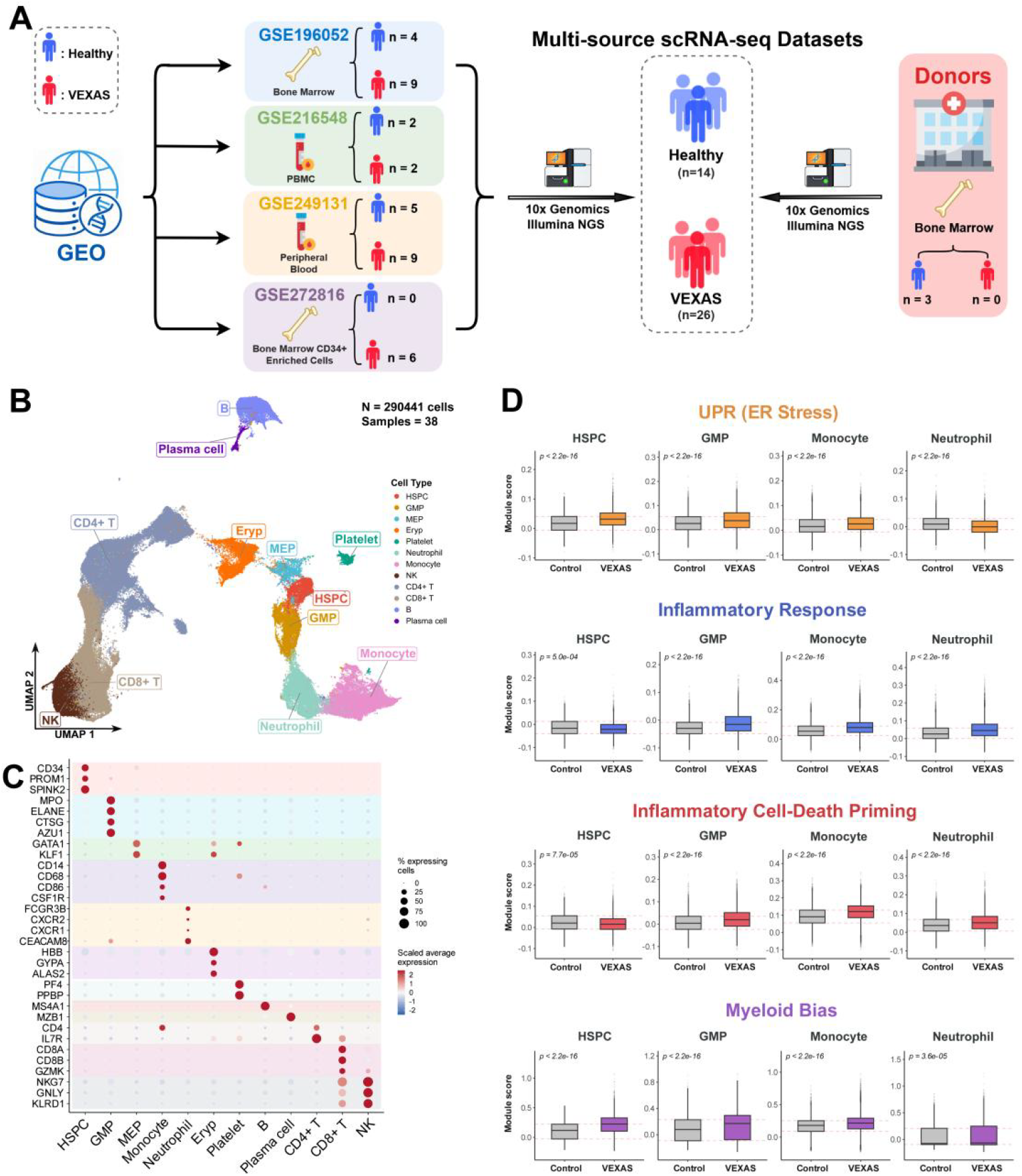
Construction of a cellular atlas of VEXAS-primed hematopoietic cells. **(A)** Study design and cohort composition. Four public GEO datasets (GSE196052, GSE216548, GSE249131, and GSE272816) and three in-house healthy bone marrow donors initially contributed 14 healthy and 26 VEXAS samples from bone marrow, CD34-enriched bone marrow, PBMCs, and whole peripheral blood. After quality control, 38 samples were retained. **(B)** UMAP of 290,441 high quality cells colored by 12 major annotated cell populations. HSPC, hematopoietic stem and progenitor cell; GMP, granulocyte-monocyte progenitor; MEP, megakaryocyte-erythroid progenitor; EryP, erythroid progenitor. **(C)** Dot plot of canonical lineage markers used to validate cell type annotation. Dot size denotes the percentage of expressing cells and color denotes scaled average expression. **(D)** Cell-level module scores for UPR/ER stress, inflammatory response, inflammatory cell-death priming, and myeloid bias in HSPCs, GMPs, monocytes, and neutrophils from control and VEXAS samples. Boxplots show the score distributions, and P values are displayed above comparisons.

Canonical marker-based annotation resolved 12 major hematopoietic and immune populations, including HSPCs, GMPs, megakaryocyte–erythroid progenitors, erythroid progenitors, platelets, neutrophils, monocytes, B cells, plasma cells, CD4^+^ T cells, CD8^+^ T cells, and natural killer cells (**Figure 3B**). The assignments were supported by canonical lineage markers, including *CD34* and *PROM1* in HSPCs, *MPO* and *ELANE* in granulocytic progenitors, *GATA1* and *KLF1* in erythroid progenitors, *CD14* and *CSF1R* in monocytes, and *FCGR3B* and *CXCR2* in neutrophils (**Figure 3C**).^44–46^ Bone marrow and peripheral blood cells occupied the same integrated manifold but retained their expected enrichment of progenitor and mature immune compartments, respectively.^46,47^ Similarly, healthy control and VEXAS-conditioned cells were represented within the same major populations, whereas the observed variation was consistent with differences in tissue source, dataset composition, and *CD34* enrichment (**Supplementary Fig. S5**). Thus, the integrated atlas captured a shared hematopoietic landscape and the batch effect is minimized.

We next focused on HSPCs, GMPs, monocytes, and neutrophils because these populations represent ordered progenitor and mature cell states along myeloid differentiation. ^35,46,47^ These four compartments are also directly implicated in VEXAS pathobiology, whereas other lineages, particularly lymphoid lineages, show limited tolerance to UBA1 mutations (e.g., UBA1 p.Met41Leu;).^1^ To characterize how disease-associated functional programs varied across these populations, we compared the activities of four predefined gene modules between VEXAS and healthy control cells: unfolded protein response/endoplasmic reticulum stress,^48^ inflammatory response, inflammatory cell death priming, and myeloid bias (**Figure 3D**). All four modules showed significant disease-associated alterations, although their relative activities differed among cell types. UPR/ER stress alterations were most evident in HSPCs and GMPs, whereas inflammatory response and cell death programs became more prominent in monocytes and neutrophils (**Figure 3D**). Myeloid bias scores were altered in both progenitor and mature myeloid populations, indicating that VEXAS associated dysregulation extended across the myeloid differentiation hierarchy rather than being confined to terminal effector cells.

Cell type resolved differential expression analysis provided complementary results at the gene level for these cell type-dependent functional patterns. In VEXAS-primed HSPCs and GMPs, coordinated expression changes were observed across multiple functional gene classes, including inflammatory signaling related genes, such as *NFKB1, IL18*, and *STAT3*. ^49,50^ ER stress and unfolded protein response genes are also enriched by CauNagi, such as *ATF6, EIF2AK3*, and *CALR*; and regulators of hematopoietic and myeloid differentiation such as *RUNX1, FLT3*, and *SPI1*. By contrast, mature myeloid populations exhibited a narrower and more specific to each cell type pattern of alteration, exemplified by *IL1B* and *EIF2AK3* in monocytes and *IFITM1* and *SPI1* in neutrophils (**Supplementary Fig. S6**). Lymphoid lineage-associated genes were relatively more highly expressed in healthy control cells, consistent with the established tendency toward myeloid-skewed hematopoiesis in VEXAS syndrome and the limited tolerance of lymphoid lineages to UBA1 mutations.^3–5^ These findings indicate that progenitor and mature myeloid populations share disease-associated alterations in inflammatory and stress-response programs while retaining distinct specific to each cell type expression patterns. Although these results provide gene-level support for the functional module analysis, expression changes alone are insufficient to establish alterations in lineage fate or transcription factor activity.

Together, the integrated atlas showed that VEXAS associated transcriptional changes occurred across multiple established hematopoietic populations rather than being confined to a distinct disease specific population. Across HSPCs, GMPs, monocytes, and neutrophils, these changes converged on common functional themes—including proteostasis, inflammation, cell death, and lineage regulation—but differed in magnitude and in the contributing genes among cell types. The recurrence of these functional programs across the predefined myeloid differentiation hierarchy, together with their cell type dependent variation, motivated us to perform the subsequent but more detailed cascade candidate regulator (CCR) analysis to identify regulators whose support was maintained, attenuated, or amplified across these populations.

### Multi-dimensional propagation scores assist to identify 36 core CCRs and a layered regulatory network

Identifying CCRs requires an initial candidate pool that is both reproducible across cell types and biologically relevant to the disease. We therefore first examined whether high priority global candidate regulators (GCR) recurred across cell types more frequently than expected under matched random backgrounds. Candidates generated from the CauNagi’s Minimum feedback vertex set (MFVS) showed greater excess recurrence than genes selected solely from the global top 5% ranking. The observed minus null recurrence was 0.61 versus 0.37 within the myeloid top 10% sets and 1.63 versus 0.93 across the all cell top-10% sets (**Supplementary Fig. S7**). We next performed pathway enrichment analysis ^11,12^ of the GCRs to determine whether the candidate pool captured biological processes relevant to VEXAS. These candidates were enriched in inflammatory and immune pathways, including TNF/NF-κB, NOD-like receptor, and interferon-γ signaling, as well as pathways related to hematopoietic differentiation, heme metabolism, ER protein processing, ubiquitin–proteasome degradation, hypoxia, apoptosis, and cell cycle regulation (**Figure 4A**). The implicated processes were consistent with major inflammatory, proteostatic, cell-death, and hematopoietic abnormalities previously reported in VEXAS syndrome.^7–10^ Together, the across cell type recurrence and functional enrichment results supported the reproducibility and biological relevance of the pool of CCRs, providing a solid rationale for subsequent cascade classification and layered network construction.

**Figure 4:**
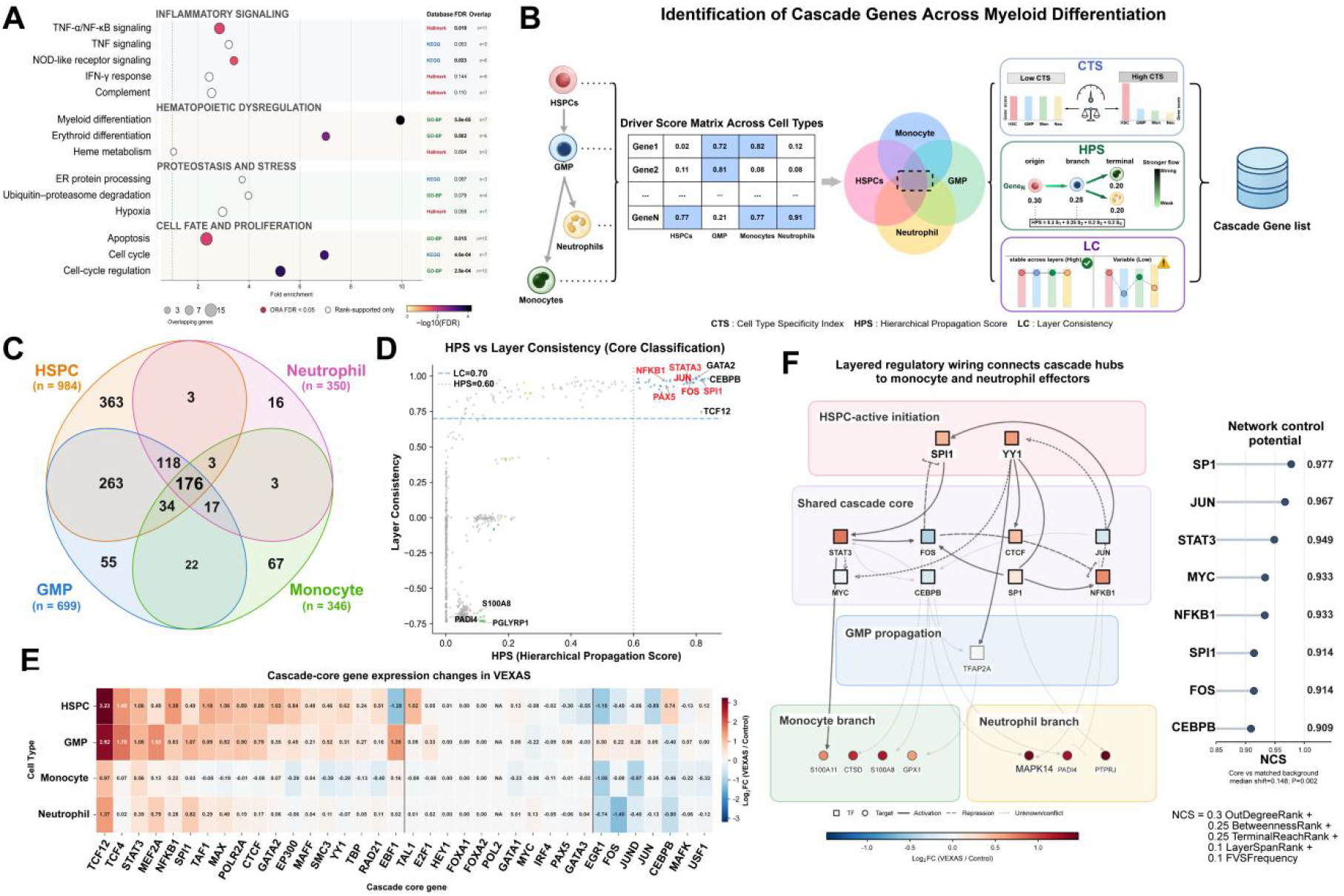
Identification of cascade candidate regulators and reconstruction of a layered VEXAS regulatory network. **(A)** Functional enrichment of CauNagi-prioritized VEXAS global candidate regulators across inflammatory signaling, hematopoietic dysregulation, proteostasis/stress, and cellfate/proliferation categories. Point position denotes fold enrichment, point size denotes the number of overlapping genes, color denotes −log10(FDR), and filled points indicate significant over-representation. **(B)** Schematic of CCRs identification. Cell type-specific driver scores from HSPC, GMP, monocyte, and neutrophil trajectories are assembled into a common matrix and summarized by CTS, HPS, and LC before cascade-role assignment. **(C)** Intersection of CCRs identified in HSPC (n = 984), GMP (n = 699), monocyte (n = 346), and neutrophil (n = 350) respective trajectories. The central overlap contains 176 genes. **(D)** HPS-versus-LC landscape used for core classification. Dashed thresholds indicate HPS = 0.60 and LC = 0.70; representative high-propagation/high-consistency genes and terminal-effector genes are labeled. **(E)** Heatmap of log2 fold changes (VEXAS versus control) for the 36 shared CCRs across HSPCs, GMPs, monocytes, and neutrophils. **(F)** Layered directed network connecting HSPC initiators, shared CCRs, GMP propagators, monocyte amplifiers, and neutrophil amplifiers. Node color denotes log2 fold change, edge style denotes activation, repression, or unresolved/conflicting direction, and the right panel reports network control scores for prioritized regulators. The core group showed a median NCS increase of 0.148 relative to matched background (P = 0.002, calculated using 10,000 matched permutations).

To distinguish shared genes from state-restricted regulatory network support, we assembled cell type-specific driver scores into a gene-by-cell-type matrix and summarized each candidate by three orthogonal measures (**Figure 4B**): (a). cell type-specific score (CTS)^51^ for quantifying the concentration of support within individual cell types; (b). the hierarchical propagation score (HPS) for summarizing the support across HSPCs, GMPs, monocytes, and neutrophils, with greater weight assigned to earlier differentiation states; and (c). layer consistency (LC) for measuring the stability of scores across these populations. Candidates with low CTS generally exhibited higher HPS and LC, whereas candidates restricted to individual cell types showed lower layer-crossing support and consistency (**Supplementary Fig. S8B**). Together, these measures characterized the breadth, hierarchical distribution, and stability of candidate support, enabling classification beyond simple overlap between gene lists.

The HSPC, GMP, monocyte, and neutrophil analyses identified 984, 699, 346, and 350 candidate genes, respectively, with 176 genes shared across all four cell type sets (**Figure 4C**). When setting a bar for requiring a non-zero driver score in every cell type, together with HPS ≥ 0.60, LC ≥ 0.70, and CTS ≤ 0.20, we finally refined this shared gene-set to 36 uniform and core CCRs. The lineage-specific tasks additionally identified 12 initiators at the level of HSPC, 13 propagators at the level of GMP, one amplifier in monocytes (*S100A8*), and three amplifiers in neutrophils (*PADI4, PGLYRP1, AGTPBP1*). We also noticed that no decay type core met the prespecified criteria. Eight CCRs and the single monocyte amplifier (*S100A8*) had prior literature support for their involvement in VEXAS-associated regulation, whereas other CCRs are new for understanding the pathology of VEXAS syndrome and should be experimentally tested in future (**Supplementary Fig. S8A**). In the joint HPS–LC distribution, candidates including *NFKB1, STAT3, CEBPB, SPI1, JUN*, and *FOS* occupied the region of high propagation and consistency, whereas *S100A8* and *PADI4* exhibited score patterns consistent with terminal effector roles (**Figure 4D**). This computational criterion distinguished candidate regulators supported by disease-related cell types at specific positions or branches of the myeloid hierarchy from those without pathological relevance.

We next examined the drivers specific to each cell type with scores and expression profiles of the CCRs to characterize their regulatory support and transcriptional changes. *SPI1, CEBPB*, the *AP-1* factors *JUN, JUND*, and *FOS*, and the inflammatory regulators *NFKB1* ^49^ and *STAT3* ^50^ received substantial driver score support across all four cell types. Other candidates showed variation across differentiation states, including stronger support for *TCF12* in early progenitors and differences between the two mature myeloid branches (**Supplementary Fig. S9**). Despite broadly elevated driver scores across cell types, the expression heatmap revealed heterogeneous VEXAS–control expression differences among these populations (**Figure 4E**). Thus, the shared regulatory support did not necessarily reflect a uniform association with disease expression pattern.

Expression analysis along the differentiation trajectories showed that *STAT3* was elevated in VEXAS syndrome along both myeloid branches, whereas *SPI1, FOS*, and *NFKB1* showed significant differences along the neutrophil branch but not the monocyte branch (**Supplementary Fig. S10**). Thus, sustained cascade support did not require consistently elevated transcript abundance across both branches, as CauNagi also incorporated target gene dynamics, transcription factor evidence, and network position.

To place the prioritized CCRs within a regulatory network, we integrated iDREM transcription-factor (TF) evidence with directed TF–target relationships from TRRUST and CollecTRI.^52,53^ Candidates were mapped to HSPC-initiation, shared-core, GMP-propagation, monocyte, or neutrophil layers, and regulatory edges were retained according to the predefined hierarchy and supporting evidence. The resulting network connected HSPC-associated regulators such as *SP1* and *YY1* with shared hubs including *STAT3, FOS, JUN, NFKB1, CEBPB*, and *SPI1*, and with branch-restricted downstream effectors (**Figure 4F**). Network control scoring (NCS) placed *SP1, JUN, STAT3, MYC, NFKB1, SPI1, FOS*, and *CEBPB* among the leading nodes. Compared with a covariate-matched background, cascade-core genes exhibited a higher median network control score, with a median difference of 0.148 (P = 0.002), indicating that they occupied comparatively prominent positions under this network-based measure.

Together, the outcomes from the CauNagi implemented in VEXAS syndrome refined the global candidate pool into 36 cascade-core genes and additional candidates with differentiation-state-specific support. Their driver scores, expression dynamics, and inferred regulatory relationships provided complementary descriptions of shared and celltype-dependent regulation across the myeloid hierarchy.

### Cascade regulators converge on VEXAS mechanisms and show conserved functional relevance

Following the identification of 36 core CCRs and their organization into a layered regulatory network, we examined the expression patterns of representative regulators and their associated downstream gene programs. We also systematically evaluated support for these candidates by independent datasets from murine VEXAS-like models and from human bulk RNA-seq. These analyses assessed whether these prioritized genes are associated with the transcriptional abnormalities observed in VEXAS syndrome and whether the selected candidates have additional support from complementary mouse and human transcriptomic data.

Based on their high priority and prominent positions in the layered regulatory network, we first selected *SPI1, NFKB1, STAT3* and *FOS*. To examine whether their downstream targets also exhibited expression changes in VEXAS syndrome, we selected four representative target genes for each transcription factor using curated regulatory relationships and compared their expression between the VEXAS-primed and control cells. In the pooled comparisons, *PRTN3, TNFAIP3, SOCS3*, and *CXCL8*, representing targets of *SPI1, NFKB1, STAT3*, and *FOS*, respectively, showed increased expression, whereas targets such as *CSF1R* and *CCL2* showed decreased expression (**Figure 5A**). These findings revealed changes in the expression of downstream targets of all four regulators, with differences in direction among individual genes. To characterize the functional relevance of these changes beyond the selected examples, we performed enrichment analysis of the target sets for each regulator in the four cell types. Enriched pathways included hematopoietic differentiation, cytokine signaling, Toll-like receptor and NOD-like receptor signaling, and JAK–STAT and PI3K–Akt signaling, with variation among cell types (**Supplementary Fig. S11**). This functional characterization defined the target programs subsequently examined for disease-associated changes.

**Figure 5:**
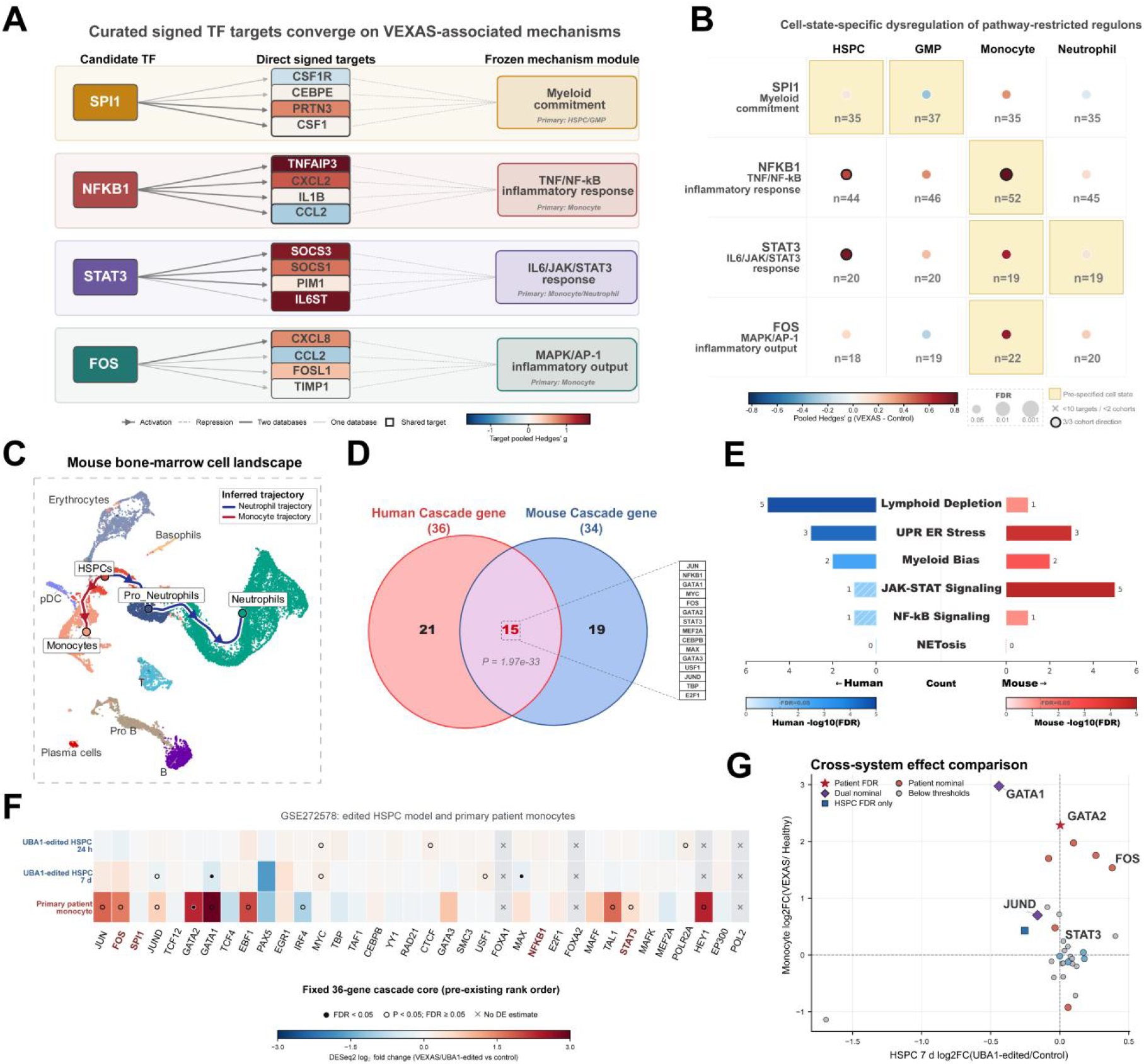
Functional convergence and external validation of VEXAS cascade regulators. **(A)** Biological interpretation of prioritized CCRs in VEXAS syndrome. Curated regulatory links between four prioritized CCRs—SPI1, NFKB1, STAT3, and FOS—and their pathway-restricted targets obtained from TRRUST and CollecTRI. Target color represents the pooled Hedges’ g for VEXAS versus control; arrow type indicates activation or repression, line thickness indicates support from one or both databases, and double borders denote shared targets. **(B)** Cell-state-specific effect profiles of pathway-restricted regulons across HSPCs, GMPs, monocytes, and neutrophils. Color represents pooled Hedges’ g, and point size represents −log_10_(BH-adjusted FDR). Highlighted boxes mark prespecified cell states, black outlines indicate concordant directions across three cohorts, crosses indicate insufficient coverage, and n denotes the number of retained targets. **(C)** Myeloid differentiation trajectory inferred from bone-marrow single-cell data from an in-house murine VEXAS model, comprising HSPCs, monocytes, Neutrophil_1, and Neutrophil_2 (OMIX006178). **(D)** Intersection of the human 36-gene and mouse 34-gene CCRs sets after mapping mouse genes to human orthologs. Fifteen CCRs were shared (one-sided hypergeometric test; background = 18,000 genes; P = 1.97×10^−33^). **(E)** Comparison of mechanism categories represented by human and mouse CCRs. Bar length indicates gene count, color intensity represents −log_10_(BH-adjusted FDR), and hatching denotes nonsignificant enrichment. **(F)** Differential-expression effect map of the fixed 36-gene cascade core in UBA1-edited HSPCs at 24 h and day 7 and in primary VEXAS monocytes relative to their respective controls (GSE272578). Color represents DESeq2 log_2_ fold change; filled circles indicate FDR < 0.05, open circles indicate (P < 0.05) with FDR ≥ 0.05, and crosses indicate unavailable estimates. HSPC P values are experiment-internal because the samples are technical replicates. **(G)** Comparison of log_2_ fold changes between day-7 UBA1-edited HSPCs and VEXAS monocytes. Red stars indicate monocyte FDR significance; purple diamonds, nominal significance in both comparisons; blue squares, HSPC FDR significance only; orange and light-blue circles, nominal significance only in monocytes and HSPCs, respectively; and gray circles, values below both thresholds. Dashed lines indicate zero change.

To determine how these programs varied across cell types, we calculated regulon scores using targets assigned to predefined pathways for each sample and cell type in three VEXAS cohorts (**Figure 5B**). *NFKB1* and *FOS* showed the strongest module increases in monocytes, while *STAT3* showed alterations across several myeloid populations and *SPI1* exhibited more moderate pooled effects. These patterns linked the four regulators to distinct downstream responses across myeloid differentiation. Analysis of the complete *NFKB1* target set further revealed differences among functional groups. Immune regulation, adhesion, and NF-κB feedback groups showed positive mean expression changes, whereas cell cycle and survival groups showed negative changes **(Supplementary Fig. S12)**. Thus, the target analyses identified variation both across cell types and among the downstream programs associated with a shared regulator.

We next assessed whether CauNagi could identify similar CCRs in an independent mouse model of VEXAS syndrome with *UBA1* dysfunction. We applied the framework to bone marrow scRNA-seq datasets from mice with conditional *UBA1* deletion in neutrophils developed in our laboratory (*S100a8Cre*;*Uba1*^*fl/y*^; OMIX006178).8 HSPCs, monocytes, and neutrophil states were used to reconstruct myeloid differentiation trajectories, followed by candidate selection using the cascade scoring strategy (**Figure 5C**). The human and mouse analyses identified 36 and 34 CCRs, respectively, with 15 shared candidates, including *JUN, NFKB1, FOS, GATA2, STAT3*, and *CEBPB*. This overlap significantly exceeded the random expectation (**Figure 5D**). Functional comparison showed that UPR/ER stress, myeloid bias, and NF-κB signaling were represented in both candidate sets, whereas JAK– STAT signaling was more strongly represented in the mouse set and genes associated with lymphoid depletion were more prominent in the human set (**Figure 5E**). These results demonstrated that CauNagi recovered a subset of the human cascade candidates in the mouse model while revealing both shared and distinct features of their functional disturbance caused by UBA1 loss of function in myeloid cells.

We next evaluated the 36 cascade core genes using independent human bulk RNA-seq data from HSPCs collected 24 hours and days 7 after *UBA1* editing and from VEXAS patient and control monocytes (GSE272578).^4^ Expression changes varied across time points and cellular backgrounds (**Figure 5F**). Comparison of day 7 HSPCs with patient monocytes showed increased *FOS* expression in both systems, a more pronounced *GATA2* increase in patient monocytes, and opposite directions of change for *GATA1* and *JUND* (**Figure 5G**). Although no core gene reached FDR significance in both comparisons, the patient monocyte candidate set was enriched for cascade core genes (FDR = 0.00013). These results provided support for the candidate set in independent human data while revealing variation in the expression responses of individual genes.

Together, these analyses from mouse models and human bulk RNA-seq datasets linked highly prioritized CCRs rather than simply GCRs to altered downstream programs, recovered a subset of the human candidates in an independent mouse analysis, and provided additional expression support from other available human bulk RNA-seq data complementary to the single-cell datasets. These findings support the relevance of selected candidates to VEXAS syndrome and help prioritize the cell types and downstream target genes to be examined in future experimental settings.

## Discussion

In this study, we reported a causal inferring tool called CauNagi, which has an iterative computational framework for identifying candidate regulators across ordered cellular states. Built on the concept-level causal disentanglement and concept conditioned diffusion modeling constructed by CausCell,^20^ as well as on the iterative coupling of cell representation, dynamic trajectory analysis, iDREM regulatory inference ^22^, and gene weight feedback introduced by UNAGI ^21^, CauNagi integrates these components into a unified and streamlined workflow for hierarchical regulator prioritization. It further evaluates candidates using cell type specificity, hierarchical propagation strength, and layer consistency, thereby distinguishing regulators with shared support across the HSPC–GMP– monocyte/neutrophil hierarchy from those restricted to particular cell states. When applied to VEXAS syndrome, CauNagi provides a developmental stage-resolved view of disease-associated regulation across myeloid differentiation. Of note, the stages analyzed here (for VEXAS syndrome) represent predefined positions within the hematopoietic hierarchy rather than longitudinal clinical progression within individual patients, but disease-stages datasets are also compatible to CauNagi when longitudinal data are available. This particular task is being reflected in the third module of CauNagi, the CCR module. This distinction reflects the structure of the VEXAS datasets used in the present study rather than an inherent limitation of CauNagi, which can also accommodate longitudinal clinical samples when such data are available.

This positioning module also distinguishes CauNagi from other regulatory analysis methods. SCENIC reconstructs regulons and uses regulon activity to characterize cellular states, but it does not iteratively update cell representations using trajectory-derived regulatory evidence.^54^ iDREM identifies branching expression programs and their associated regulators but operates downstream of a supplied temporal structure.^22^ CEFCON and WMDS.net prioritize regulators through lineage specific network inference or network control analysis without jointly refining the underlying representation.^17,18^ CauFinder combines causal disentanglement with network control for a specified cell state or phenotype transition,^34^ whereas CauNagi is designed to integrate evidence across multiple ordered and branching cellular states and to assign candidates to putative hierarchical roles. When compared with UNAGI, which was developed primarily for analyzing disease progression dynamics, CauNagi focuses on cell type-crossing regulatory continuity, cascade role classification, and layered regulatory network construction.^21^

As an initial benchmark, we evaluated two core capabilities of CauNagi using independent disease datasets. In the benchmark with IPF datasets, CauNagi achieved the highest ARI and NMI among the compared representation learning methods, indicating that its latent representation effectively preserved biologically meaningful cell state structure. The ablation analysis showed that removing causal disentanglement impaired cell state separation more substantially than disabling iterative gene weight feedback, highlighting the central role of the causal disentanglement module. In the benchmark with AML datasets, CauNagi outperformed the compared methods in the top-k ranking evaluation, and its top 20 globally prioritized regulators were enriched in AML-associated pathways. Together with the IPF benchmark results, these findings support the quality of the latent representations and candidate regulator pool that underpin the subsequent cascade analysis.

When applied to the multi-source VEXAS atlas, another major part of the study, CauNagi distinguished candidate regulators with broadly sustained or cell type restricted support across HSPCs, GMPs, monocytes, and neutrophils. Using a predefined HSPC–GMP– monocyte/neutrophil hierarchy, rather than treating these compartments as temporal stages of clinical progression, the framework assessed how regulatory evidence was maintained, attenuated, or reorganized across myeloid differentiation. Collectively, these findings support a model in which related regulators may connect early hematopoietic abnormalities with mature myeloid inflammatory programs through mechanistically distinct, dependent on cellular context processes, without implying a single pathway transmitted linearly across the myeloid hierarchy.

Mouse and human transcriptomic analyses provided additional support for selected candidates. Partial overlap between human and *UBA1*-deficient mouse cascade genes indicated that a subset of prioritized regulators was reproducible across model systems.^8^ Human bulk RNA-seq from *UBA1*-edited HSPCs and VEXAS patient monocytes provided complementary evidence that several candidates or their associated programs were dysregulated in both early and mature hematopoietic compartments.^4^ Together with the trajectory-resolved and target program analyses, these observations support the disease relevance of selected candidates and define specific regulators and downstream programs for experimental testing.

### Limitations

Although CauNagi effectively disentangled biological concepts from residual variation and enabled the inference of candidate key regulators from the VEXAS single-cell atlas, several limitations should be considered. Given the limited availability of the VEXAS scRNA-seq datasets and samples, the VEXAS atlas generated in this study integrates datasets derived from different sample types (e.g., BMMCs and PBMCs), cohorts, and sequencing protocols. Although batch correction reduced dataset- and sample-associated variation, the reconstructed hierarchy should be interpreted as cross-sectional regulatory continuity rather than direct longitudinal, clonal, or clinical progression. VEXAS syndrome is a clonal hematopoiesis-driven disease; however, for sample-level analyses, we assumed that all VEXAS-primed cells from the patient cohort carried UBA1 mutations, although the reported mutant chimerism is approximately 70 – 90%.^55^ The learned representation also depends on the prespecified concept graph and supplied annotations, while network construction is constrained by the coverage and context specificity of existing TF–target databases.^52,53^ Finally, although prior biological information was incorporated into the framework, CauNagi identifies computationally supported regulatory hypotheses rather than experimentally established causal effects. Thus, CauNagi may assist in generating deeper biological hypotheses. Targeted perturbation experiments,^56^ clone-resolved longitudinal sampling, single-cell multi-omics, and systematic parameter-sensitivity analyses will therefore be required in the future to evaluate the predicted regulatory roles. Furthermore, application of CauNagi to additional disease scenarios with ordered or branching cellular transitions (i.e., disease evolutions between clonal hematopoiesis (CH), low-risk of MDS, high-risk of MDS and AML during myeloid malignancies; or MGUS, SMM and MM during lymphoid malignancies) will further strengthen its generalizability and robustness.

## Conclusions

Here we introduce a new and powerful computational framework CauNagi as a general useful working pipeline for identifying candidate regulators that may sustain, propagate, or amplify disease-associated programs across ordered cellular states and for generating testable hypotheses about regulatory dynamics. We benchmarked the tool and it outperformed several competing methods. We further applied CauNagi to VEXAS syndrome, a recently recognized hematologic and autoinflammatory disorder, and prioritized several candidate key regulators, a subset of which was supported by independent mouse-model datasets. CauNagi will be applicable to other progressive disorders for which multistage single-cell transcriptomic datasets are available.

## Abbreviations

VEXAS Diseases and Hematopoiesis related

AML: Acute myeloid leukemia
BM: Bone Marrow
ER: Endoplasmic reticulum
EryP: Erythroid progenitor
ETF: Effective transcription factor
GMP: Granulocyte-monocyte progenitor
HSPC: Hematopoietic stem and progenitor cell
IPF: Idiopathic pulmonary fibrosis
MDS: Myelodysplastic syndrome
MEP: Megakaryocyte-erythroid progenitor
MGUS: Monoclonal gammopathy of undetermined significance
MM: Multiple myeloma
NK: Natural killer cell
NOD: Nucleotide-binding oligomerization domain, used in NOD-like receptor
PBMC: Peripheral blood mononuclear cell
SMM: Smoldering multiple myeloma
TF: Transcription factor
UPR: Unfolded protein response

## Abbreviations

CauNagi Machine Learning and Computations related

ARI: Adjusted Rand index
ASW: Average silhouette width
CCRs: Candidate cascade regulators
cDAG: Concept-level causal directed acyclic graph
cLISI: Cell type local inverse Simpson’s index
CTS: Cell type specificity index, a measure of support concentration across cell types
DEG: Differentially expressed gene
F1: F1 score; the harmonic mean of precision and recall
FVS: Feedback Vertex Set
GCR: Global candidate regulators
GEO: Gene Expression Omnibus
GSEA: Gene Set Enrichment Analysis
HPS: Hierarchical propagation score, a measure of support propagation across the cellular hierarchy
LC: Layer consistency, a measure of stability across cell type layers
NCS: Network control scores, a measure of relative regulatory influence in the network
NMI: Normalized mutual information
OD: Out-degree
PCA: Principal Component Analysis
PR: PageRank
PE: Pathway enrichment
RNA-seq: RNA sequencing
UMAP: Uniform Manifold Approximation and Projection; an embedding and visualization method
UMI: Unique molecular identifier

## Methods

### CauNagi framework and input representation

CauNagi was developed for single-cell transcriptomic datasets containing predefined disease stages or ordered time points. Each stage was stored as an AnnData object^57^ with a common gene feature space and cell level annotations. Data from all stages were merged, and disease stage, cell type, and available covariates were encoded as discrete biological concepts. A further unexplained concept dimension was included to capture variation not represented by the supplied labels; accordingly, the user provided concept causal adjacency matrix contained one more dimension than the number of observable concepts^20^. Disease stages were defined from the study design and were not inferred as pseudotime within the model. Expression values were normalized and log transformed when required. When the number of genes exceeded the training limit, the highest ranked highly variable genes were retained.

### Concept-conditioned generative diffusion module

A Gaussian diffusion model ^20,58^ was used to learn a noise-robust representation of the single-cell expression distribution. During the forward process, Gaussian noise was added to the original expression vector x0 according to a linear noise schedule:

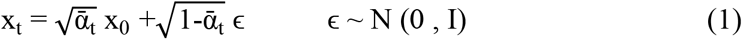

where t is a randomly sampled diffusion time step, alpha_t is the cumulative noise schedule parameter, and epsilon follows a standard multivariate normal distribution. The denoising network received the noisy expression vector, a sinusoidal time embedding, and the disentangled concept representation and was trained to reconstruct x0. L2 reconstruction loss was used by default, with L1 loss available as an alternative. Cross attention fused expression and concept embeddings so that generation was conditioned on disease stage, cell identity, and other supplied concepts. The attention operation modeled conditional dependence between expression and concept representations and was not interpreted as a gene-gene attention network.

### Causal disentanglement module

The encoder predicted the mean and log variance of an exogenous latent representation from each expression vector. Exogenous variables were sampled by reparameterization:

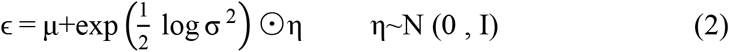

A fixed concept-level causal adjacency matrix A was then used to propagate exogenous disturbances into endogenous concept representations:

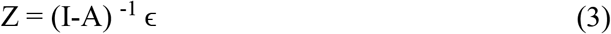

Each known concept corresponded to an independent latent subspace, whereas the unexplained subspace captured residual expression variation. The concept causal graph was specified from prior biological knowledge and remained fixed during training. Concept classification losses encouraged each known subspace to predict its corresponding observed label. An adversarial loss reduced the ability of the unexplained subspace to predict known concepts, thereby limiting information leakage, and a Kullback-Leibler divergence regularized the exogenous representation toward a standard normal prior. The total loss was:

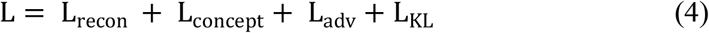

where L_recon, L_concept, L_adv, and L_KL denote reconstruction, supervised concept, adversarial, and latent prior losses, respectively.^20^

### Within stage cell state identification

After training, the cell type concept subspace was used for state analysis. Multiple latent samples were averaged for each cell to reduce sampling variability. Local cosine distances were used to estimate representation uncertainty and to construct latent distribution parameters compatible with across stages matching. For each disease stage, a k-nearestneighbor graph was built in the latent space, Leiden clustering^59^ was performed, and PAGA^60^ was used to summarize state connectivity and initialize UMAP.^61^ A clustering parameter optimization (CPO) module adjusted the number of neighbors and Leiden resolution separately for each stage to improve comparability across datasets with different cell numbers or heterogeneity. For every cluster, CauNagi calculated mean expression, the top 100 cluster marker genes, cell type composition, and latent distribution summaries.

### Across stages state graph and dynamic trajectories

Clusters from adjacent disease stages were compared pairwise using the symmetric Kullback-Leibler divergence between their latent distributions and the expression distance between their top 100 marker genes. Each distance type was min-max normalized and the two components were summed to produce a composite state distance. For every cluster in the later stage, the nearest cluster in the preceding stage was treated as the candidate source. A cumulative probability derived from the composite distance distribution was used to retain supported links. Directed edges across all adjacent stages were then combined into a state transition graph spanning all disease stages.

### iDREM based dynamic regulatory analysis

For each trajectory connecting all disease stages, cluster level mean expression was ordered by stage and supplied to iDREM ^22^ together with species-specific TF-target priors. Effective transcription factors were retained according to iDREM significance. Genes that increased or decreased between adjacent stages were ranked by the absolute expression change, and high confidence transcription factors and dynamic targets were matched to human or mouse TF-target priors. This procedure generated trajectory and stage specific candidate regulatory relationships informed by dynamic expression, transcription factor activity, and prior directionality.

### Iterative gene weight feedback

iDREM transcription factors, dynamic target genes, and stage-dependent expression changes were converted into cluster specific gene weights. Genes with strong dynamic changes and supporting TF-target evidence received greater weights; transcription factors received an additional increment. At each iteration, the previous weight was decayed and combined with newly acquired evidence:

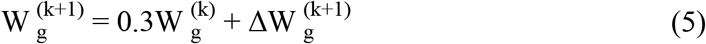

The updated geneWeight layer was used to weight gene wise reconstruction errors in the next training round. This created an iterative loop of representation learning, state clustering, trajectory construction, iDREM inference, weight updating, and retraining.

### Dynamic marker identification and candidate set construction

After the final iteration, Wilcoxon tests were used to identify genes associated with the hierarchy of latent state clusters, with multiple testing correction applied across genes. In parallel, dynamic markers were defined as genes showing sustained increases or decreases along iDREM trajectories. Their trajectory associated statistics were compared with random background distributions and adjusted for multiple testing. Genes supported by at least one of four CauNagi outputs (nonzero final iterative gene weights, significant iDREM effective transcription factors, significant dynamic markers, or hierarchical cluster markers) were retained as the candidate pool for downstream prioritization. This step defined the candidate set and its evidence features; numerical integration and global ranking were performed as described below.

### Global multi evidence scoring and driver gene ranking

For each gene in the candidate pool defined above, a composite score was calculated as a weighted sum of four continuous evidence components and one binary hierarchical marker indicator. Genes were then ranked in descending order of their composite scores for global prioritization. First, the iterative gene weight score combined the mean final geneWeight across stages with a penalty for cross-stage variability:

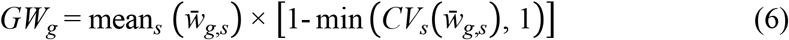

Second, iDREM evidence was defined by the maximum effective-transcription-factor significance across nodes, weighted by the number of trajectories supporting that regulator:

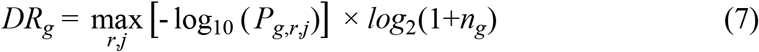

Third, dynamic-marker evidence integrated statistical significance, effect magnitude, and cross-trajectory recurrence using the minimum adjusted P value, maximum absolute log2 fold change, and number of supporting trajectories:

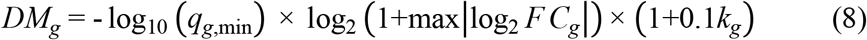

Fourth, network-topology evidence was derived from a directed disease-associated regulatory network constructed from iDREM transcription factors, dynamic markers, genes with nonzero final geneWeight, and prior TF-target edges. Normalized betweenness centrality (B), PageRank (PR), out-degree (OD), and minimum feedback vertex set membership were integrated as follows ^62–64^:

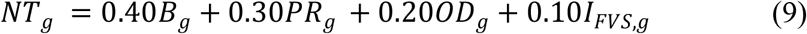

Fifth, hierarchical-cluster-marker support was represented by a binary indicator:

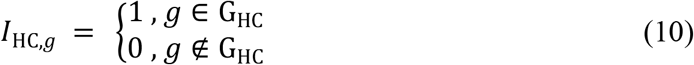

For each gene, iterative gene weight, iDREM regulatory evidence, dynamic-marker support, and network-topology evidence were separately min–max normalized across the candidate gene set and combined as a weighted sum to obtain its base driver score:

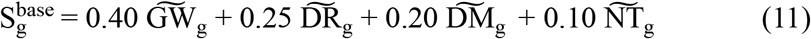

Finally, the base driver score was adjusted using the hierarchical-cluster-marker indicator, a gene class correction factor Cg, and a multiplier rewarding concordant support from multiple evidence sources to calculate the global driver score for each gene:

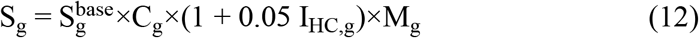

The gene class correction moderately upweighted transcription factors and downweighted hemoglobin, ribosomal, immunoglobulin, and mitochondrial genes to limit rankings driven primarily by expression abundance or cell-composition effects. The final score was interpreted as a relative prioritization of disease-associated regulatory potential, not as a probability of causality.

### Cell type mapping and cell type-specific driver scores

Final-iteration Leiden clusters were mapped to annotated cell types. A cell type occupying at least 50% of a cluster was assigned as the dominant identity; otherwise, the two most abundant identities were recorded as a mixed annotation. iDREM trajectory filenames were parsed to map starting and terminal clusters to cell types, and only trajectories assigned to one cell type were retained for cell type-specific analyses. Within cell type c, the maximum iDREM and dynamic-marker evidence scores for gene g were normalized, and its cell typespecific driver score was calculated using the following formula:

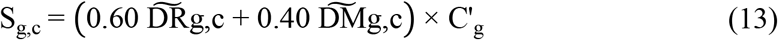

where C’_g is the gene-class correction. Genes were ranked separately in HSPC, GMP, monocyte, and neutrophil trajectories.

### Cascade-gene quantification and classification

The union of HSPC, GMP, monocyte, and neutrophil candidate lists was assembled into a gene-by-cell type driver matrix. Duplicate gene entries within a cell type were represented by their maximum score, and genes not detected in a cell type were assigned zero. To preserve relative differences across the four cell types, scores were normalized using the global minimum and maximum:

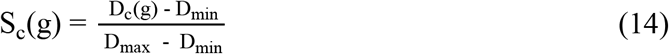

For each gene, normalized scores S_HSPC, S_GMP, S_Mono, and S_Neut were converted to proportional contributions p_c(g):

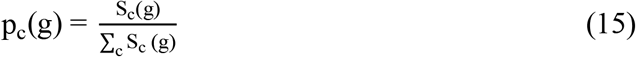

cell type specificity was defined using normalized Shannon entropy^51^:

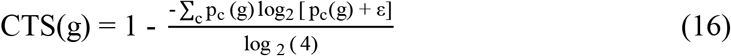

with ε= 10 ^-12^ to avoid logarithms of zero. CTS approaches zero for uniform cross-cell support and one for cell type-restricted support.

Hierarchical propagation strength weighted upstream compartments more heavily:

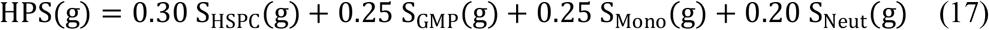

Layer consistency was calculated by subtracting the coefficient of variation of the normalized driver scores for each gene across the four cell types from one:

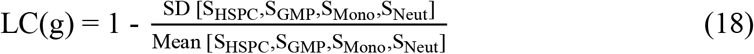

LC was set to zero when the four-layer mean was zero. Propagation depth was the number of cell types with a raw driver score above 0.01. Uniform cascade-core genes were required to satisfy:

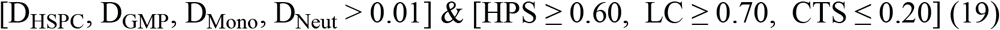

Genes with support in all four layers and high HPS but a progressive decline from HSPC to terminal cells were classified as decay type cascade genes. Remaining genes were assigned, in fixed priority order, as HSPC initiators, GMP propagators, monocyte amplifiers, or neutrophil amplifiers. Four-set overlap significance was evaluated with 10,000 permutations. If N_obs is the observed four-way intersection and N_b is the intersection in permutation b, the empirical P value was:

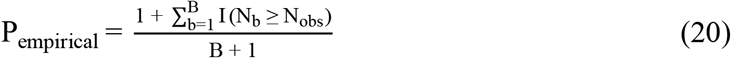

### Layered regulatory network and network control score

Effective transcription factors, node significance, and node frequency were extracted from final-iteration iDREM output files (DREM.json) ^22^ for the four myeloid trajectories. Records with ETF P values below 0.05 were retained. Within each trajectory, significance and node-frequency measures were percentile normalized and integrated as:

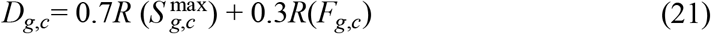

where Dg,c denotes the iDREM evidence score for gene g in the trajectory corresponding to cell type c, Sg,c is the maximum node-level significance score, Fg,c is the node-occurrence frequency, and R denotes percentile normalization within the trajectory.

Directed TF-target relationships from TRRUST and CollecTRI ^52,53^ were harmonized by gene symbol while preserving direction, database source, and PMID evidence. Relationships supported by both resources were assigned a database-support value of 1.00; single-database edges received 0.70; conflicting directions were flagged rather than forced. Candidate genes were mapped to HSPC initiation, shared core, GMP propagation, monocyte, or neutrophil layers according to their cascade classification.

An edge was retained when both nodes belonged to non-Other cascade categories, the source TF had significant iDREM activity in the relevant trajectory, at least one regulatory database supported the edge, the target had a nonzero layer score, the direction was compatible with the prespecified hierarchy, and the edge was not self-regulatory. Permitted edges respected the directional layer structure and excluded lateral links between the monocyte and neutrophil terminal branches. Edge scores were calculated as:

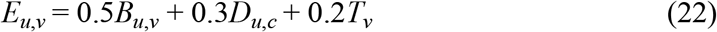

where Eu,v denotes the score of the directed edge from source TF (u) to target gene (v), Bu,v is the database-support weight, Du,c is the iDREM evidence score of the source TF in the relevant trajectory, and Tv is the target gene score in its assigned layer.

Tier 1 edges were supported by both databases and Tier 2 edges by one database. For visualization, edges with stronger support, higher scores, cross-layer connectivity, or terminal-node reach were prioritized; each target was limited to two incoming edges and each regulator to four outgoing edges. Network control scores (NCS) combined ranked out-degree, betweenness, terminal reach, layer span, and feedback-vertex-set frequency ^62–64^:

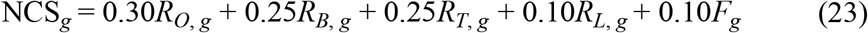

Feedback-vertex-set frequency was estimated by edge resampling. A null NCS distribution for cascade-core genes was generated from 10,000 random sets matched for TF identity, iDREM detection, number of active cell types, mean expression, and prior network outdegree.

### VEXAS single-cell datasets and preprocessing

Four public scRNA-seq datasets were collected and combined with data from three inhouse healthy bone marrow samples. GSE196052 and GSE272816 provided bone marrow data, with CD34+ progenitor enrichment in GSE272816, whereas GSE216548 and GSE249131 provided peripheral blood data^4,5,7,9^. For datasets requiring read-level processing, SRA files were downloaded using SRA Toolkit^65^, converted to FASTQ using fasterq-dump, and assessed using FastQC^66^. Cell Ranger^67^ was used for sequence alignment and gene expression quantification against the GRCh38.p14 reference genome with matched GENCODE annotations. Sample specific gene-by-cell expression matrices were obtained for subsequent quality control and integration. Expression matrices or previously saved R objects were imported into Seurat 4.3.0 ^68^ in R 4.2.0, with sample identifier, dataset, tissue source, and disease status incorporated into the cell metadata. Quality control applied dataset specific thresholds for the number of detected genes and total UMI counts, and mitochondrial and hemoglobin transcript percentages. Samples PT9 and bm2.BM were excluded because of abnormal quality profiles, including elevated hemoglobin-transcript fractions, and predicted doublets were removed using DoubletFinder^42^. The retained data were merged and normalized, followed by selection of 3,000 highly variable genes, data scaling, and principal component analysis (PCA) with 50 components. Dataset- and sample-associated batch effects were corrected using Harmony ^43^. The first 40 Harmony components were used for graph construction and UMAP visualization with 35 neighbors, and Leiden clustering was performed at a resolution of 0.5. Cluster markers were identified using the FindAllMarkers function (Wilcoxon rank-sum test), and cell types were manually annotated using CellMarker ^44^, PanglaoDB ^45^, canonical markers, and published single-cell studies. Following the exclusion of suspected mixed-cell clusters, these procedures yielded an integrated VEXAS single-cell atlas comprising 12 major cell types.

### Cross-species validation

Bone marrow scRNA-seq data from a neutrophil-specific Uba1 conditional knockout mouse model (*S100a8Cre*;*Uba1*^*fl/y*^,OMIX006178)^8^ were processed and analyzed using the same preprocessing strategy and CauNagi analysis workflow. The HSPC, Monocyte, Neutrophil_1, and Neutrophil_2 cell populations were used to reconstruct the mouse myeloid trajectory. Mouse candidate cascade genes were mapped to human ortholog symbols before cross-species gene set comparison. The observed human-mouse overlap was evaluated against a null distribution generated by permutation, and disease related functional categories were compared between species using the same curated functional definitions.

### External validation with human bulk RNA-seq

GSE272578 ^4^ comprised 19 bulk RNA-seq samples: *UBA1*-edited and control HSPCs collected 24 h after editing (three and two samples, respectively), *UBA1*-edited and control HSPCs collected at day 7 (three and three samples), and monocytes from five patients with VEXAS and three healthy donors. Differential expression analysis was performed for each experimental comparison with multiple testing correction, yielding respective lists of differentially expressed genes for downstream integration. Effect directions of the 36 cascade core genes were compared across edited HSPCs and patient monocytes. Gene set enrichment analysis (GSEA) ^69^ was performed to determine whether disease-associated bulk RNA-seq candidates were overrepresented among the cascade core genes.

### Ablation study

To assess the contributions of causal disentanglement and iterative gene weight feedback to the representation-learning performance of CauNagi, we conducted ablation studies on the cell embedding task using the IPF dataset (GSE286182)^21^. The full CauNagi model was compared with two ablated variants, designated w/o causal disentanglement and w/o iterative feedback. In the w/o causal disentanglement variant, the concept-level causal transformation and the associated supervised and adversarial disentanglement constraints were removed, while the generative backbone was retained. In the w/o iterative feedback variant, feedback of iDREM derived gene weights to subsequent training rounds was disabled. As in the representation learning benchmark, each configuration was run independently ten times with different random seeds and evaluated using the same scIB metrics^30^.

## Data availability

The datasets analyzed or generated in this study are available as described below. The four publicly available single-cell RNA-sequencing datasets used to construct the integrated VEXAS atlas are available in the Gene Expression Omnibus (GEO) under accession codes GSE196052[https://www.ncbi.nlm.nih.gov/geo/query/acc.cgi?acc=GSE196052], GSE216548[https://www.ncbi.nlm.nih.gov/geo/query/acc.cgi?acc=GSE216548], GSE249131[https://www.ncbi.nlm.nih.gov/geo/query/acc.cgi?acc=GSE249131], and GSE272816[https://www.ncbi.nlm.nih.gov/geo/query/acc.cgi?acc=GSE272816]. The idiopathic pulmonary fibrosis and acute myeloid leukemia datasets used for method benchmarking are available in GEO under accession codes GSE286182[https://www.ncbi.nlm.nih.gov/geo/query/acc.cgi?acc=GSE286182] and GSE116256[https://www.ncbi.nlm.nih.gov/geo/query/acc.cgi?acc=GSE116256], respectively. The independent human bulk RNA-sequencing dataset used for external validation is available in GEO under accession code GSE272578[https://www.ncbi.nlm.nih.gov/geo/query/acc.cgi?acc=GSE272578]. The mouse bone marrow single-cell RNA-sequencing data from the neutrophil-specific Uba1 conditional-knockout model used for cross-species validation are publicly available in the National Genomics Data Center OMIX repository under accession code OMIX006178 [https://ngdc.cncb.ac.cn/omix/release/OMIX006178].

The prior transcription factor–target interactions used for regulatory-network construction were obtained from TRRUST v2[https://www.grnpedia.org/trrust/] and CollecTRI[https://github.com/saezlab/CollecTRI], with CollecTRI regulons accessed through OmniPath. The AML reference gene set assembled from DepMap, DisGeNET, published experimental evidence, and expert review is provided in the Supplementary Information. All other data supporting the findings of this study are available within the article and its Supplementary Information.

## Code availability

The CauNagi algorithm is implemented in Python. The source code of CauNagi is available at https://github.com/steamed-stuffed-bun/CauNagi.

## Acknowledgments

We thank our colleagues for technical support, critically reading our manuscript, and their suggestions to improve the manuscript.

## Author Contributions

DB designed the study, performed experiments, analyzed data, and wrote the first draft of the manuscript. HY, JL, GD, and SX performed experiments and analyzed data. ZC conceived and designed the study, supervised the study, and critically revised the manuscript for important intellectual content. All authors approved the final version of the manuscript. ZC served as the guarantor of the study.

