## Supplementary Materials for "Inferring cascade drivers of VEXAS syndrome by a causal machine learning tool CauNagi"

### Supplementary Notes

#### Supplementary Note 1: Computational resources and training time

**Hardware and training configuration.** CauNagi was applied to the VEXAS atlas containing 290,441 cells represented by 3,000 selected genes. CauNagi was run on a single NVIDIA GeForce RTX 3090 GPU with 24 GiB of device memory, with the mini-batch size set to 64. The outer CauNagi procedure was repeated six times. For each iteration, epoch_num was set to 3,000 (the training log records indices 0–3,000), max_profile_size was set to 4,000, and model checkpoints and samples were saved every 1,500 epochs. Two disease stages were specified, with disease stage and cell type supplied as observed concepts to the model.

**GPU memory requirement and runtime.** During the VEXAS run, the GPU used for computation occupied 18,254 MiB (17.83 GiB) of device memory. Therefore, a GPU with 24 GiB of memory was used for the present configuration. According to the timestamped training records, the complete six-iteration workflow required approximately 7 h 23 min (7.4 h). Of this time, the six neural-network fitting stages together accounted for approximately 45 min, whereas most of the remaining elapsed time occurred between fitting stages, during which CauNagi mainly performed iterative cell-state analysis, regulatory network reconstruction, and gene-weight feedback. Therefore, under the same settings, approximately 7–10 h should be reserved on an RTX 3090-class GPU for an analysis of a dataset comparable in size to the VEXAS atlas. This estimate does not include additional post-run analyses that were not timestamped in the training log.

**CPU memory guidance.** The geneWeight layer used in this analysis had dimensions of 290,441 × 3,000. When stored as float32, one dense matrix of this size occupies approximately 3.25 GiB of memory, and an expression matrix of the same dimensions requires an additional 3.25 GiB. CPU execution must also accommodate normalized or scaled data copies, model and optimizer tensors, latent representations, neighbour graphs, AnnData metadata, and temporary objects. For the present VEXAS configuration, we therefore recommend at least 64 GiB of host RAM, with 128 GiB preferred to reduce the risk of paging during CPU-only execution. These values are capacity recommendations rather than measured CPU peak memory. CPU-only execution is expected to require substantially longer runtime than the RTX 3090 GPU run.

**Scaling to new datasets.** For N cells and G genes, the storage required by one dense float32 cell-by-gene matrix is:

$$M_{matrix} \approx \frac{4NG}{2^{30}}\, GiB \quad \text{(S1)}$$

At 3,000 genes, the raw storage required by a single dense float32 cell-by-gene matrix is approximately 1.12 GiB for 100,000 cells, 3.25 GiB for 290,441 cells, and 11.18 GiB for 1,000,000 cells. During analysis, multiple full-size matrices and additional objects, including normalized data, geneWeight, model tensors, latent representations, graph structures, and temporary variables, may coexist in memory. Therefore, as a practical guideline, approximately 32 GiB of RAM is recommended for datasets of up to about 100,000 cells, 64–128 GiB for datasets containing several hundred thousand cells, and 128–256 GiB for datasets approaching one million cells, assuming approximately 3,000 genes.

With max_profile_size fixed at 4,000 and batch size at 64, neural-network fitting memory is partly constrained, while full expression, `geneWeight`, graph, and downstream objects continue to scale with cell number. A 24-GiB GPU was validated for the present configuration; smaller GPUs may require a reduced batch size and additional testing. Based on the VEXAS run, approximate runtimes on an RTX 3090-class GPU are 2–3 h for 50,000 cells, 3–5 h for 100,000 cells, 5–8 h for 200,000 cells, 7–10 h for ~290,000 cells, 12–18 h for 500,000 cells, and 24–36 h for 1,000,000 cells. Only the 290,441-cell run was measured; all other values are planning estimates and may vary with dataset complexity.

**Supplementary Note 2: Benchmarking cell representations**

**Dataset and comparison design.** Representation quality was evaluated using the idiopathic pulmonary fibrosis (IPF) dataset GSE286182. After preprocessing, the final AnnData object contained 27,647 cells and 2,484 genes. CauNagi[1] was compared with UNAGI (v0.5.1)[2], scGNN (v1.0.3)[3], scGGAN[4], GraphSCC[5], scGen (v2.1.0)[6], scvi-tools (v1.4.0)[7], scGPT (v0.2.4)[8] and Geneformer (v0.1.0)[9]. CauNagi, scGGAN and GraphSCC did not have formal versioned releases and are therefore reported without version numbers. Each method was run independently ten times, and the resulting cell embeddings and evaluation metrics were collected for comparison.

**Evaluation inputs and software.** The published IPF cell-type annotations were used as the common biological reference. For every method, its learned representation was stored in adata.obsm[embed_key] and evaluated with the same function. The reference cell-type labels, method-specific cluster assignments and disease-stage labels were read from adata.obs[cell_type_key], adata.obs[cluster_key] and adata.obs[batch_key], respectively. In the recorded analysis these fields corresponded to name.simple, a method-specific clustering column such as UNAGI, and stage, while the evaluated embedding was stored under a key such as z. The implementation used scIB v1.1.7[10] together with Scanpy[11] and scikit-learn[12].

**Metric implementation.** Adjusted Rand index (ARI), normalized mutual information (NMI), Davies-Bouldin index (DBI), neighbourhood label agreement, cluster-based silhouette width, cell-type silhouette width, isolated-label ASW, isolated-label F1 and graph cLISI were calculated separately for each run. The following definitions and parameter settings reproduce the supplied evaluation function.

**Adjusted Rand index.**[13] ARI was calculated as **adjusted_rand_score** (adata.obs[cell_type_key], adata.obs[cluster_key]). For the contingency table between the reference labels U and cluster assignments V, let n_ij be the number of cells shared by reference class i and cluster j, and let a_i and b_j be the corresponding marginal totals. With these quantities, ARI is:

$$A_2 = \sum_{i,j} \binom{n_{ij}}{2}$$

$$B_2 = \sum_{i} \binom{a_i}{2}$$

$$C_2 = \sum_{j} \binom{b_j}{2}$$

$$P_2 = \binom{n}{2}$$

$$ARI = \frac{A_2 - \frac{B_2 C_2}{P_2}}{0.5(B_2 + C_2) - \frac{B_2 C_2}{P_2}} \qquad \text{(S2)}$$

ARI equals 1 for identical partitions, is approximately 0 for chance-level agreement and can be negative when agreement is below chance. No additional scaling was applied.

**Normalized mutual information.** NMI was calculated as **normalized_mutual_info_score** (adata.obs[cell_type_key], adata.obs[cluster_key]). Because average_method was not supplied, the scikit-learn default arithmetic normalization was used:

$$NMI = \frac{2I(U;V)}{H(U) + H(V)} \qquad \text{(S3)}$$

Here, I(U;V) denotes mutual information and H(U) and H(V) are the entropies of the reference and cluster labels. NMI ranges from 0 to 1, with 1 indicating identical label partitions; unlike ARI, it is not adjusted for chance.

**Davies-Bouldin index.**[14] DBI was calculated as davies_bouldin_score(adata.obsm[embed_key], adata.obs[cluster_key]) using the scikit-learn defaults. It compares within-cluster dispersion with between-centroid separation in the embedding; lower values indicate more compact and better separated inferred clusters. DBI was reported as a diagnostic metric but was not included in Overall SCIB.

**Neighbourhood label score.** A 50-nearest-neighbour graph was constructed directly from the embedding with sklearn.neighbors.kneighbors_graph using mode='distance', include_self=True and n_jobs=25. For each cell, the cell-type labels of the 50 stored neighbours were compared with the focal cell's reference label. Label score was the total number of matches divided by the total number of stored neighbour relations:

$$\text{Label score} = \frac{\sum_i \left(\sum_{j \in N_{50}(i)} \delta_{y_j, y_i}\right)}{\sum_i |N_{50}(i)|} \quad \text{(S4)}$$

Although distances determined which neighbours were selected, the final agreement was unweighted; the self-neighbour was included. This study-specific measure is therefore a 50-neighbour label-agreement rate rather than a standard scIB metric.

**Silhouette scores.**[15] Two raw scikit-learn silhouette scores were calculated from the same embedding with the default Euclidean metric. Silhouette (cluster) used adata.obs[cluster_key] as the grouping, whereas Silhouette (cell type) used adata.obs[cell_type_key]. For a cell i, a(i) is its mean distance to other cells in the same group and b(i) is the smallest mean distance to another group; the cell-level silhouette and its average are:

$$s(i) = \frac{b(i) - a(i)}{max\{a(i), b(i)\}}, \; ASW = \frac{1}{n}\sum_{i=1}^{n} s(i) \quad \text{(S5)}$$

Both reported values are unscaled and range from -1 to 1. Thus, Silhouette (cluster) quantifies separation of inferred clusters, while Silhouette (cell type) quantifies separation of reference cell types. The latter is the cell-type ASW used in Overall SCIB.

**Isolated-label scores.** Isolated-label ASW and F1 were calculated with **scib.metrics.isolated_labels_asw** and **scib.metrics.isolated_labels_f1** using label_key=cell_type_key, batch_key=batch_key and embed=embed_key. No iso_threshold was supplied; under the scIB v1.1.7[10] default, isolated labels were the cell types present in the minimum number of disease stages. Isolated-label ASW contrasted each isolated cell type with all other cells in the embedding, used the default scale=True and averaged the scaled ASW values across isolated labels. For isolated-label F1, no clustering key or resolution list was supplied. scIB therefore used the default iso_label clustering and searched ten Leiden resolutions from 0.2 to 2.0 in increments of 0.2, retaining the clustering that maximized recovery of each isolated label before averaging the label-level F1 scores.

**Graph cLISI.** Before cLISI evaluation, **sc.pp.neighbors** was called with use_rep=embed_key and n_neighbors=50. The subsequent call was **scib.metrics.clisi_graph**(adata, label_key=cell_type_key, type_='embed', use_rep=embed_key). In scIB v1.1.7[10], type_='embed' recomputes a 15-neighbour graph from the specified embedding in a copy of the AnnData object, so the preceding 50-neighbour Scanpy graph does not set the cLISI neighbourhood. The default cLISI procedure retrieves k0=90 shortest-path neighbours, uses a default perplexity of 30, applies scale=True and returns the median across cells. If L_i is the raw local inverse Simpson index and C is the number of reference cell types, the purity-oriented transformation is:

$$L_i = \frac{1}{\sum_{c=1}^{C} p_{ic}{}^2}, \ cLISI_{scaled} = \frac{C - L_i}{C - 1} \quad \text{(S6)}$$

A raw LISI value near 1 denotes a locally pure cell-type neighbourhood. After transformation, values near 1 indicate stronger preservation of local cell-type identity. The batch_key argument was not passed to clisi_graph and is not used for this cell-type metric.

**Overall score and repeated runs.** The function returned all nine diagnostic entries, but Overall SCIB was calculated from exactly six values: Overall SCIB = (ARI + NMI + Silhouette (cell type) + Isolated F1 + Isolated ASW + graph cLISI) / 6. DBI, Label score and Silhouette (cluster) were excluded from this composite. The six-component score was calculated within each run; means and standard deviations shown in Supplementary Fig. S1 summarize the ten independent runs.

**Supplementary Note 3: Benchmarking candidate regulator identification**

**AML data and benchmark design.** Candidate regulator identification was evaluated using the acute myeloid leukemia (AML) single-cell dataset GSE116256 [16], processed using the same procedure described in Supplementary Note 2. CauNagi was compared with CauFinder, CEFCON, CellOracle and WMDS.net using the same preprocessed expression matrix and gene-symbol feature space. A fixed set of 36 AML-associated genes was used as the common reference set for all methods (Supplementary Table 2).

**DepMap evidence.** DepMap evidence was derived from CRISPRGeneEffect.csv[17], Model.csv and CommonEssentialControls.csv. AML models were selected from Model.csv using OncotreeLineage == 'AML'. For each gene, the AML mean gene-effect value, AML dependency fraction and AML-specific delta were calculated, and genes were retained if they met the following criteria: AML mean effect < -0.5, AML dependency fraction >= 0.5 and AML-specific delta < -0.1. Common essential genes were subsequently removed using CommonEssentialControls.csv, and the remaining candidates were restricted to transcription factors and chromatin regulators. Genes passing these filtering criteria were considered supported by DepMap evidence.

**DisGeNET and literature evidence.** The DisGeNET[18] component retained AML-associated genes with an association score > 0.9 and excluded entries supported primarily by the evidence categories Causal Mutation or Genetic Variation. The retained record does not specify a DisGeNET release identifier. A third evidence source comprised 23 AML-related publications. Candidates from DepMap, DisGeNET and the literature were integrated and reviewed to define the final reference set.

**Reference-gene evidence.** Gene-level evidence sources and supporting publications for each gene are provided in Supplementary Table 2.

**Method-specific gene scoring and ranking.** In each run, candidate genes were ranked according to the gene scores generated by each method. Because the comparison methods primarily return global candidate drivers across disease progression, whereas CauNagi focuses on candidate regulators specific to individual cell types, we introduced a Global candidate gene strategy in CauNagi to ensure comparability and fairness across methods (see Methods). This strategy integrates candidate regulators identified across all cell types and generates an overall ranking of global candidate genes based on an integrated candidate-regulator score. The score combines multiple sources of evidence, including iterative gene weights, iDREM-supported transcription factors[19], dynamic markers and regulatory-network information. CauFinder[20] ranked genes according to normalized absolute SHAP-derived causal weights. CEFCON[21] first constructed a lineage-specific regulatory network based on a prior interaction network, identified candidate driver genes supported by MFVS and MDS, and then ranked the retained regulators according to their influence_score. CellOracle[22] reconstructed regulatory networks and simulated transcription-factor knockout effects to obtain Perturbation_Score, which was subsequently used to rank candidate genes; for the AML benchmark, only genes with Perturbation_Score > 6.0 were retained. WMDS.net[23] identified candidate driver genes from a differential co-expression network using a weighted minimum-dominating-set approach, in which node weights integrate network connectivity and differential co-expression information, and candidate genes were ranked according to the gene scores generated by the method.

**Precision at k.** Let G denote the fixed 36-gene reference set and P_mr(k) the first k distinct genes in the score-descending list from method m and run r. Precision at k was evaluated for k = 1,...,20 as the proportion of the first k candidates contained in G, and the displayed curve reports the mean across ten runs:

$$Mean\ Precision_m@k = \frac{1}{10}\sum_{r=1}^{10} \frac{|P_{mr}(k) \cap G|}{k} \qquad (S7)$$

The benchmark was designed to compare the ability of different methods to score and rank candidate genes. Therefore, the relative ordering of genes within each method was evaluated, whereas the absolute magnitudes of the raw scores were not compared across methods.

**Precision, recall and F1.** For a run-specific candidate set P, the implementation converted the returned list to a set and calculated TP = |P intersect G|, FP = |P| - TP and FN = |G| - TP. Precision, recall and F1 were then calculated as:

$$Precision = \frac{TP}{TP + FP},\ Recall = \frac{TP}{TP + FN},\ F1 = \frac{2TP}{2TP + FP + FN} \quad \text{(S8)}$$

**Run-level aggregation.** Precision, recall and F1 were first calculated within each of the ten independent runs. Results were then grouped by method, and pandas mean and standard deviation were calculated for each metric. The error bars in Supplementary Fig. S3 therefore represent between-run standard deviations. F1 was averaged from run-specific F1 values rather than reconstructed from the mean precision and mean recall.

**Consensus candidate intersections.** For the UpSet[24] analysis, a method-specific consensus set was defined by counting gene occurrences across the ten runs and retaining genes present in at least five runs. The fixed 36-gene reference set was added as a separate Ground Truth set. The union of all consensus and reference genes was converted to set-membership combinations, and intersection sizes were plotted with upsetplot. Thus, Supplementary Fig. S2 summarizes reproducible candidate-set overlap across runs rather than the output of one arbitrarily selected run.

**KEGG pathway annotation.** Pathway interpretation was performed using GSEApy[25] to retrieve the KEGG_2021_Human[26] gene sets from the Enrichr[27] library. The script matched seven prespecified KEGG pathways: Acute myeloid leukemia, Hematopoietic cell lineage, Transcriptional misregulation in cancer, MAPK signaling pathway, PI3K-Akt signaling pathway, JAK-STAT signaling pathway and Cell cycle. The corresponding gene sets were then extracted and used to annotate candidate genes according to pathway membership and functional relevance.

#### Supplementary Note 4: Candidate scoring and interpretation

CauNagi-derived candidates were evaluated using complementary evidence from the final geneWeight layers[1], significant iDREM transcription factors[19], dynamic expression markers, network topology and hierarchical cell-state markers. Iterative gene-weight evidence combines the average weight across stages with its stability. iDREM evidence incorporates node-level significance and support across trajectories. Dynamic-marker evidence incorporates adjusted significance, expression-change magnitude and recurrence across trajectories. Network evidence summarizes betweenness, PageRank, out-degree and membership in a minimum feedback vertex set. A minimum feedback vertex set contains the smallest number of nodes whose removal breaks all directed cycles in the evaluated network; membership supplies a topology-based prioritization criterion.

The evidence components are normalized before weighted combination, with additional adjustments for gene categories and support from multiple evidence sources. The resulting score represents relative regulatory priority, rather than a calibrated probability of causality. Global scoring and cell-type-specific scoring use different evidence combinations and should be interpreted within their respective analyses. For the cell-type-specific analysis, trajectories are assigned using cluster cell-type composition, and trajectories involving mixed cell types are excluded. iDREM and dynamic-marker evidence from the retained trajectories then support cell-type-specific rankings.

The concept graph used during representation learning, the inferred connections between cell clusters and the TF–target regulatory network operate at distinct levels[1,2,19]. Concept-level causal constraints do not establish experimental gene-level causality, and similarity-based connections between stage-specific cell populations do not provide lineage-tracing evidence. In particular, disease-stage trajectories are distinct from the prespecified HSPC–GMP–monocyte/neutrophil hierarchy used to organize the VEXAS cascade analysis. Candidate cascade regulators are hypotheses supported by the combined computational evidence and require independent perturbation or lineage-based validation to establish causal propagation.

### Supplementary Figures

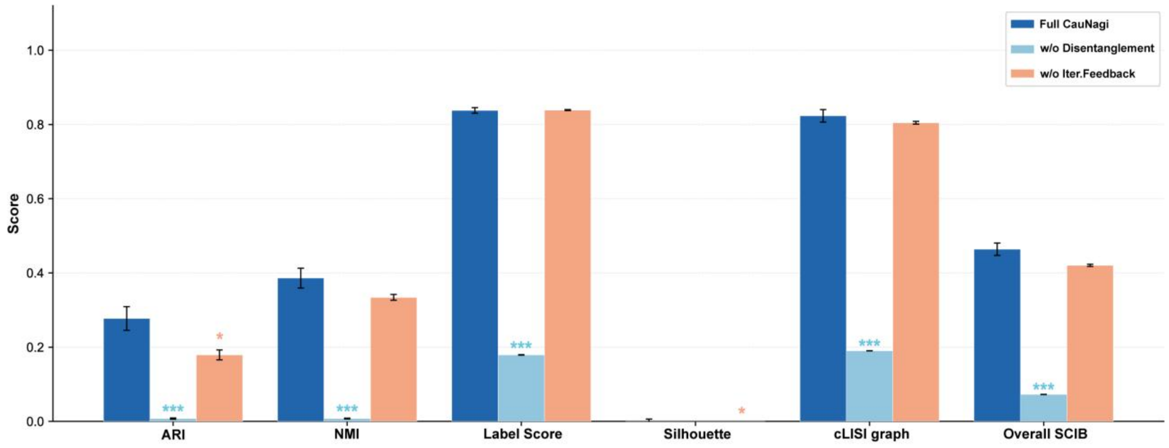

**Supplementary Figure S1: Ablation analysis of CauNagi representation learning.**

Comparison of the complete CauNagi model with variants lacking causal disentanglement or iterative gene weight feedback in the IPF dataset GSE286182. Bars show mean ARI, NMI, label score, silhouette score, graph cLISI, and overall scIB score across ten independent runs; error bars summarize between-run variation. Asterisks denote the significance of each ablated model relative to the complete CauNagi model.

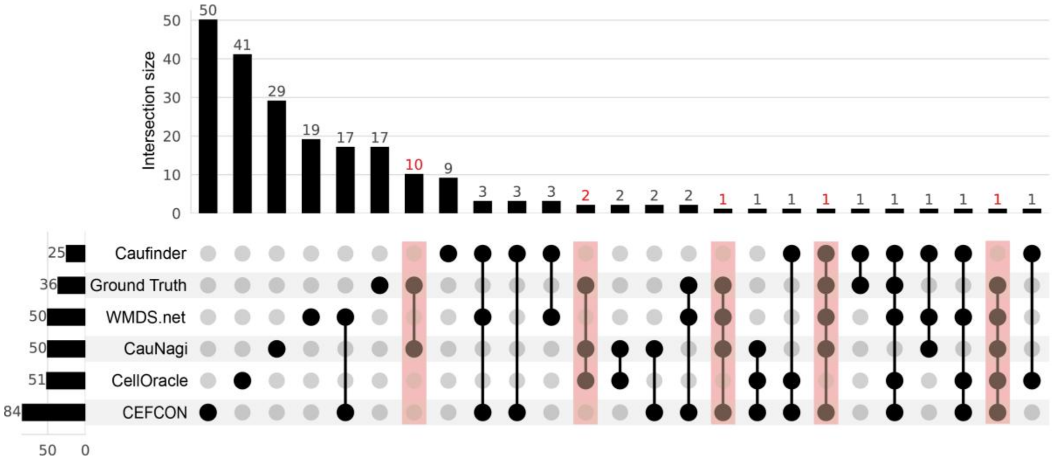

**Supplementary Figure S2: Overlap of AML candidate regulators across methods and the curated reference set.**

UpSet plot showing candidate-set intersections among CauFinder, the 36-gene AML reference set, WMDS.net, CauNagi, CellOracle, and CEFCON. Horizontal bars show total set sizes, vertical bars show intersection sizes, and the connected-dot matrix identifies the sets contributing to each intersection. Highlighted intersections contain genes from the curated AML reference set.

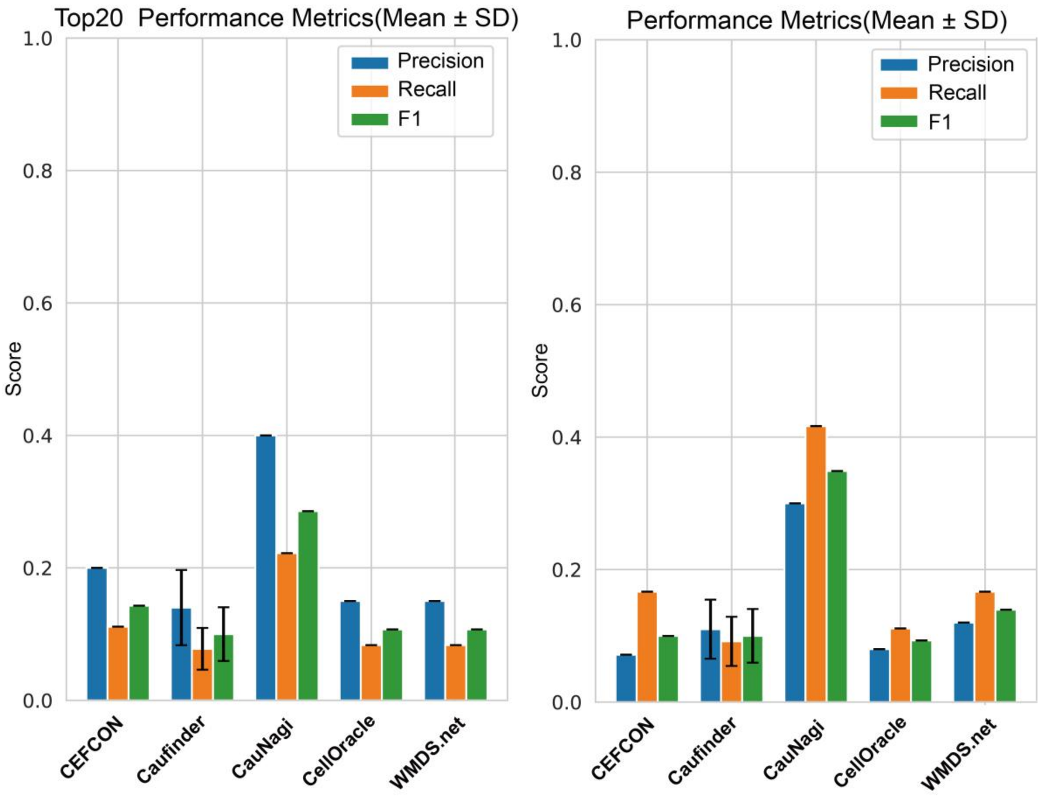

**Supplementary Figure S3: Precision, recall, and F1 performance in the AML regulator benchmark.**

Mean precision, recall, and F1 values with between-run standard deviations for CEFCON, CauFinder, CauNagi, CellOracle, and WMDS.net across ten runs. The left panel evaluates the top 20 predictions, and the right panel evaluates all reported candidates.

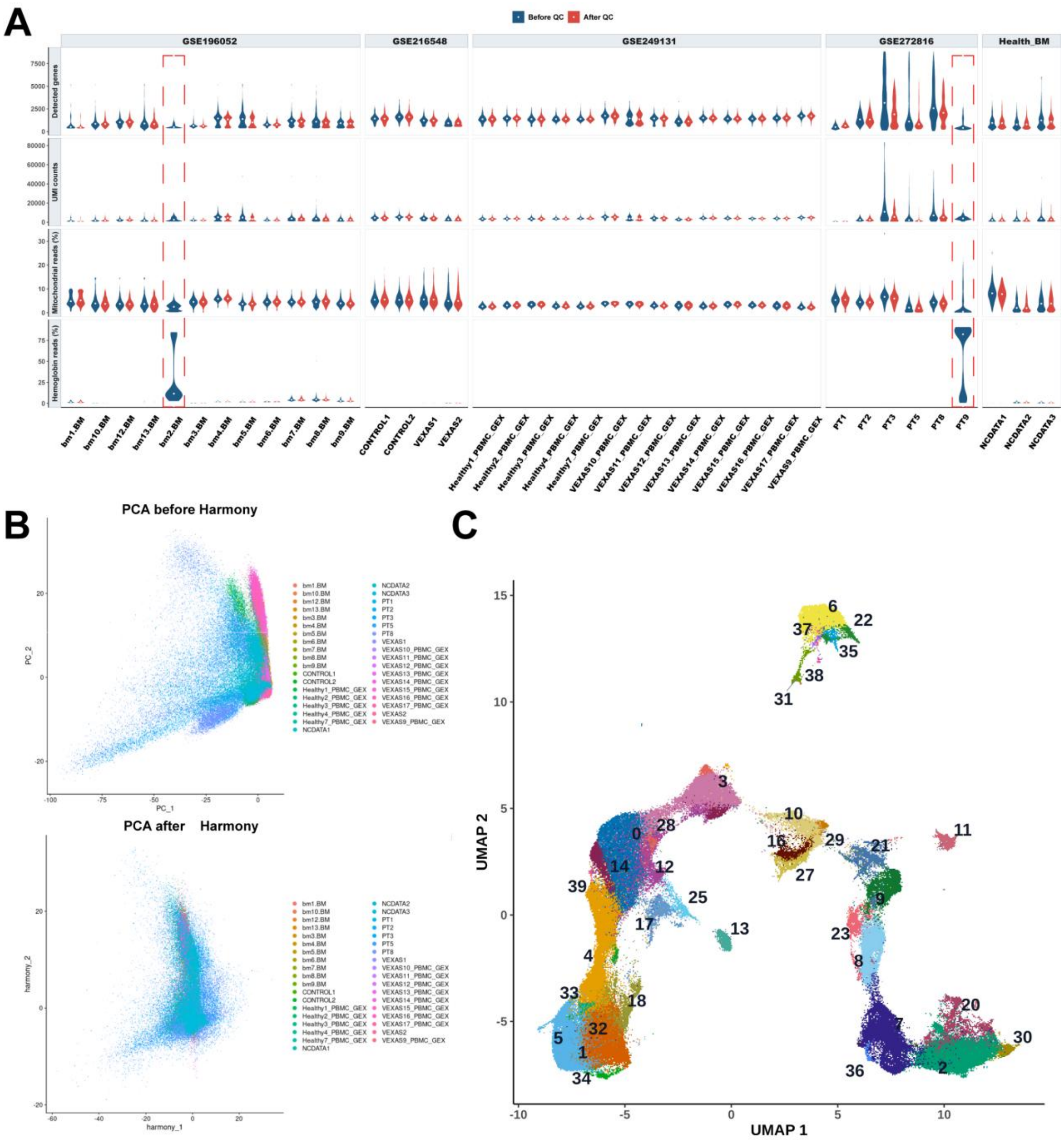

**Supplementary Figure S4: Quality control, batch correction, and initial clustering of the integrated VEXAS atlas.**

**(A)** Per-sample distributions of detected genes, UMI counts, mitochondrial-read percentage, and hemoglobin-read percentage before and after quality control. Red boxes identify the hemoglobin-contaminated samples removed from analysis. **(B)** Principal-component representations before and after Harmony correction, colored by sample. **(C)** UMAP of the Harmony-corrected data showing the 40 initial unsupervised clusters used for marker-guided annotation.

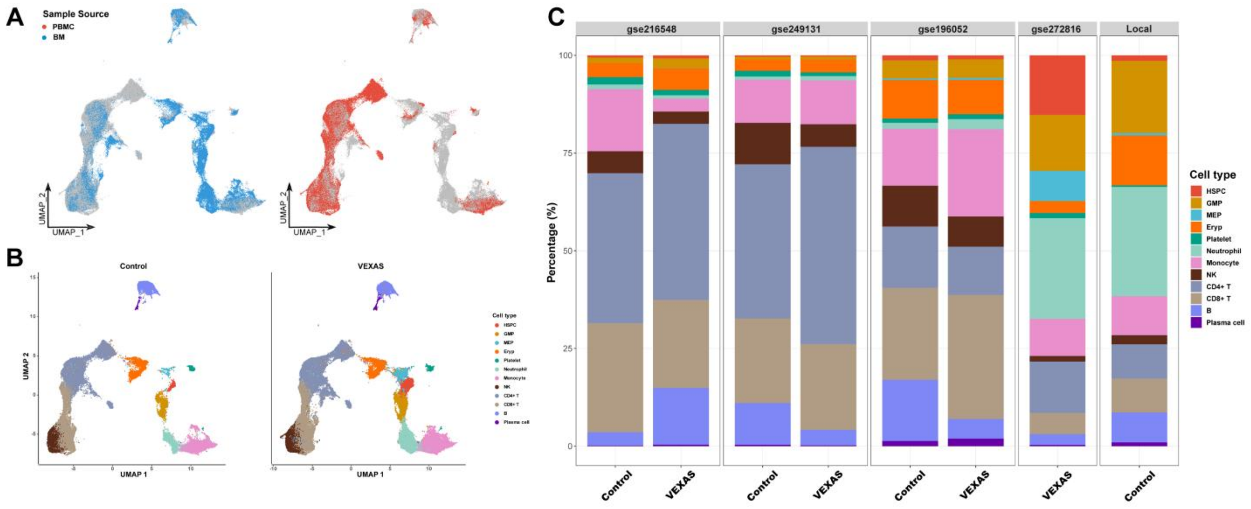

**Supplementary Figure S5: Distribution of tissue sources, disease states, and cell type composition in the VEXAS atlas.**

**(A)** UMAP projections highlighting cells derived from bone marrow and peripheral blood. **(B)** UMAP projections separated by control and VEXAS status and colored by annotated cell type. **(C)** Cell-type proportions within each dataset and disease-status group.

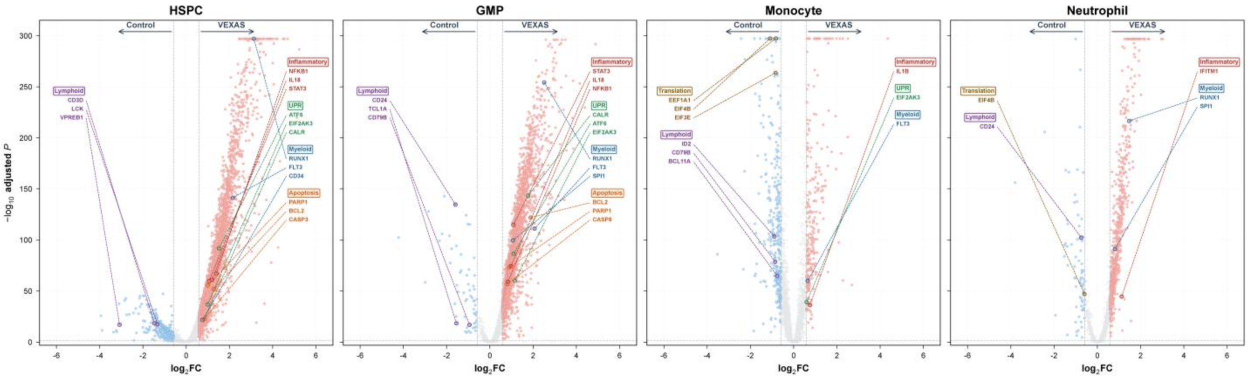

**Supplementary Figure S6: Cell type-resolved differential expression in VEXAS myeloid populations.**

Volcano plots comparing VEXAS with control cells in HSPCs, GMPs, monocytes, and neutrophils. Representative inflammatory, UPR/ER-stress, myeloid, apoptosis, lymphoid, and translation-associated genes are labeled. Point position denotes log2 fold change and adjusted statistical significance.

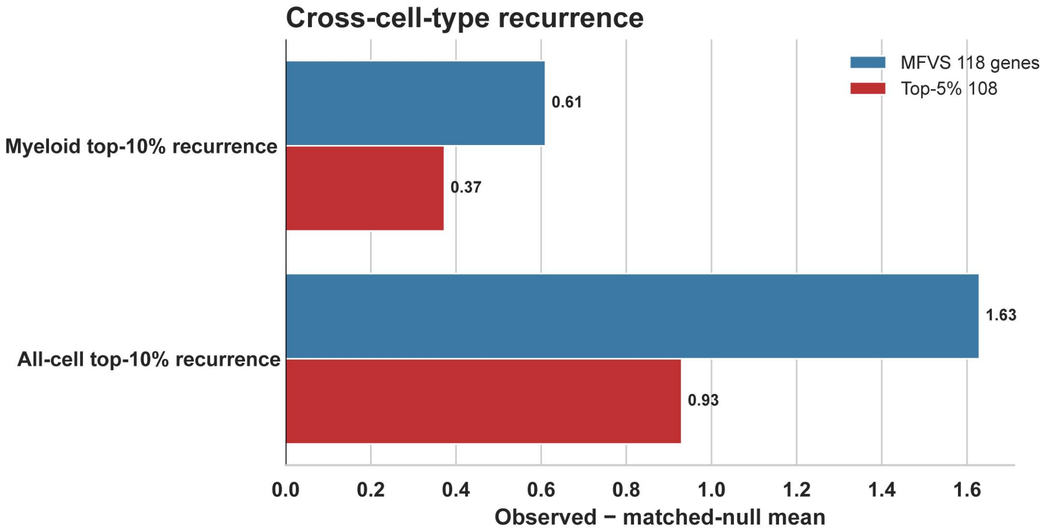

**Supplementary Figure S7: Cross-cell type recurrence of high-priority VEXAS candidate regulators.**

Observed recurrence minus the mean of covariate-matched null distributions for 118 minimum-feedback-vertex-set candidates and 108 genes in the global top 5% ranking. Recurrence was evaluated within myeloid top-10% candidate sets and across all-cell top-10% candidate sets.

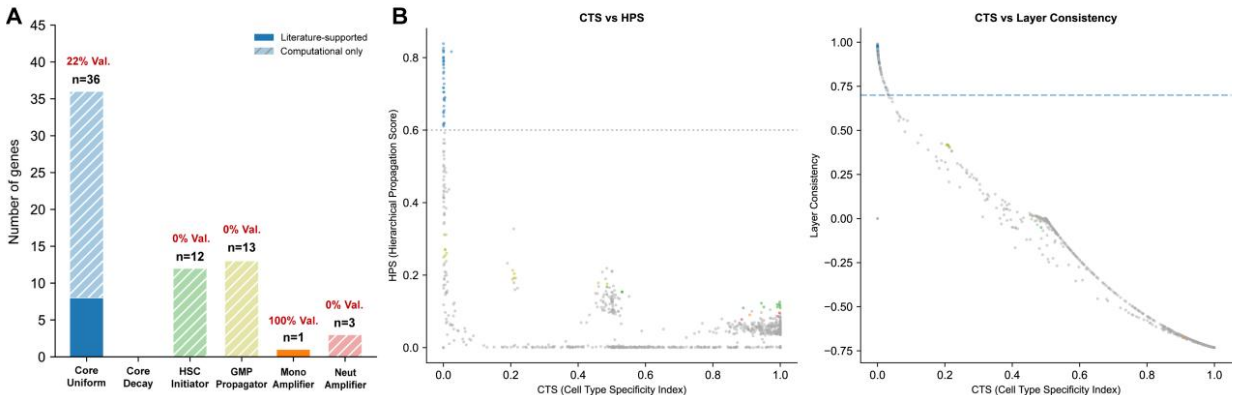

**Supplementary Figure S8: Cascade-role composition and relationships among cascade metrics.**

**(A)** Numbers of genes assigned to uniform core, decay core, HSPC initiator, GMP propagator, monocyte amplifier, and neutrophil amplifier classes. Solid segments indicate literature-supported genes and hatched segments indicate computational-only candidates; red labels report the supported percentage. **(B)** Relationships of CTS with HPS and LC. Dashed lines indicate the HPS and LC thresholds used for uniform cascade-core classification.

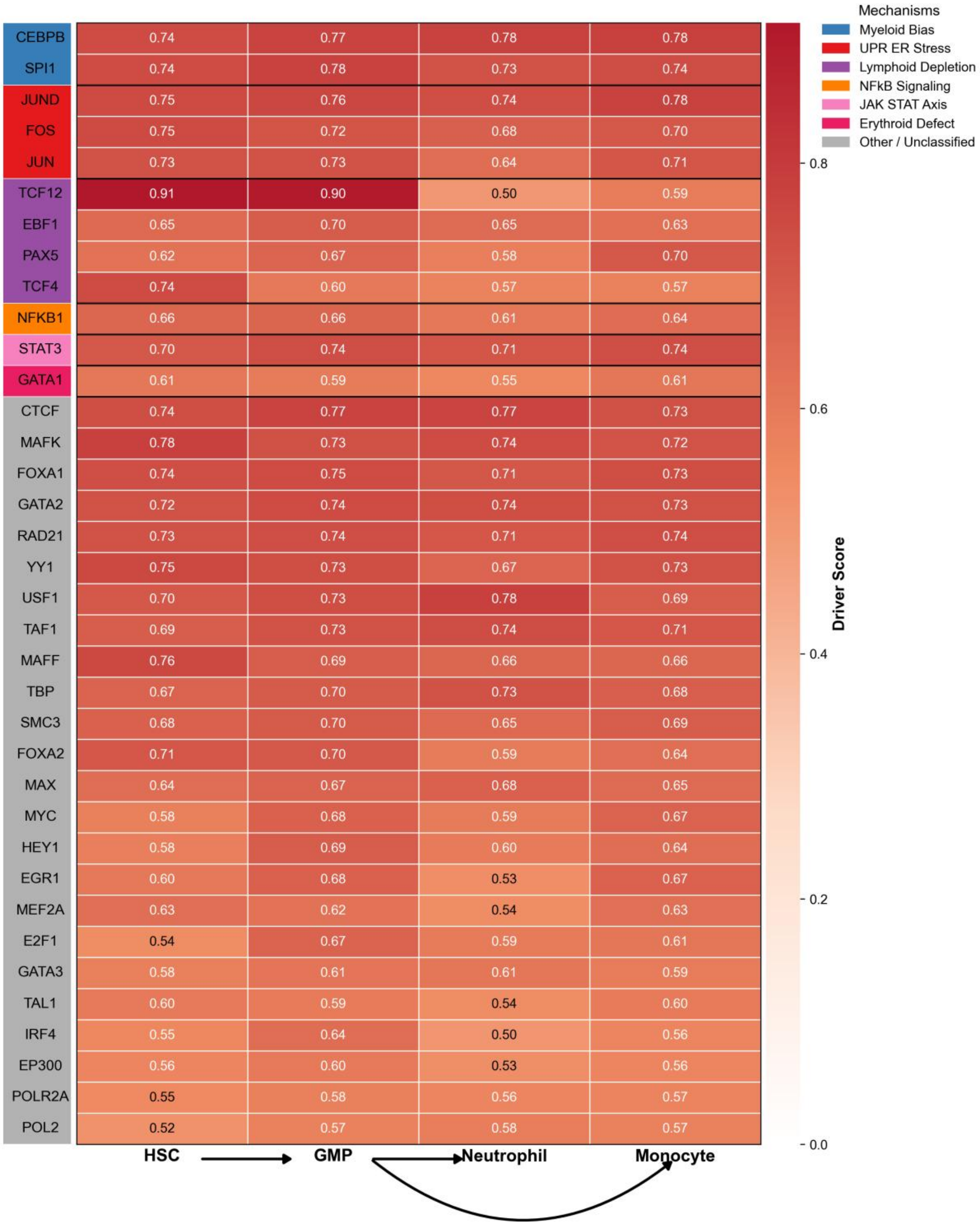

**Supplementary Figure S9: Cell type-specific driver scores of the 36 VEXAS cascade-core genes.**

Heatmap of normalized driver scores across HSPCs, GMPs, neutrophils, and monocytes. Row-side colors group genes by selected VEXAS-associated mechanisms, and arrows indicate the prespecified myeloid differentiation hierarchy.

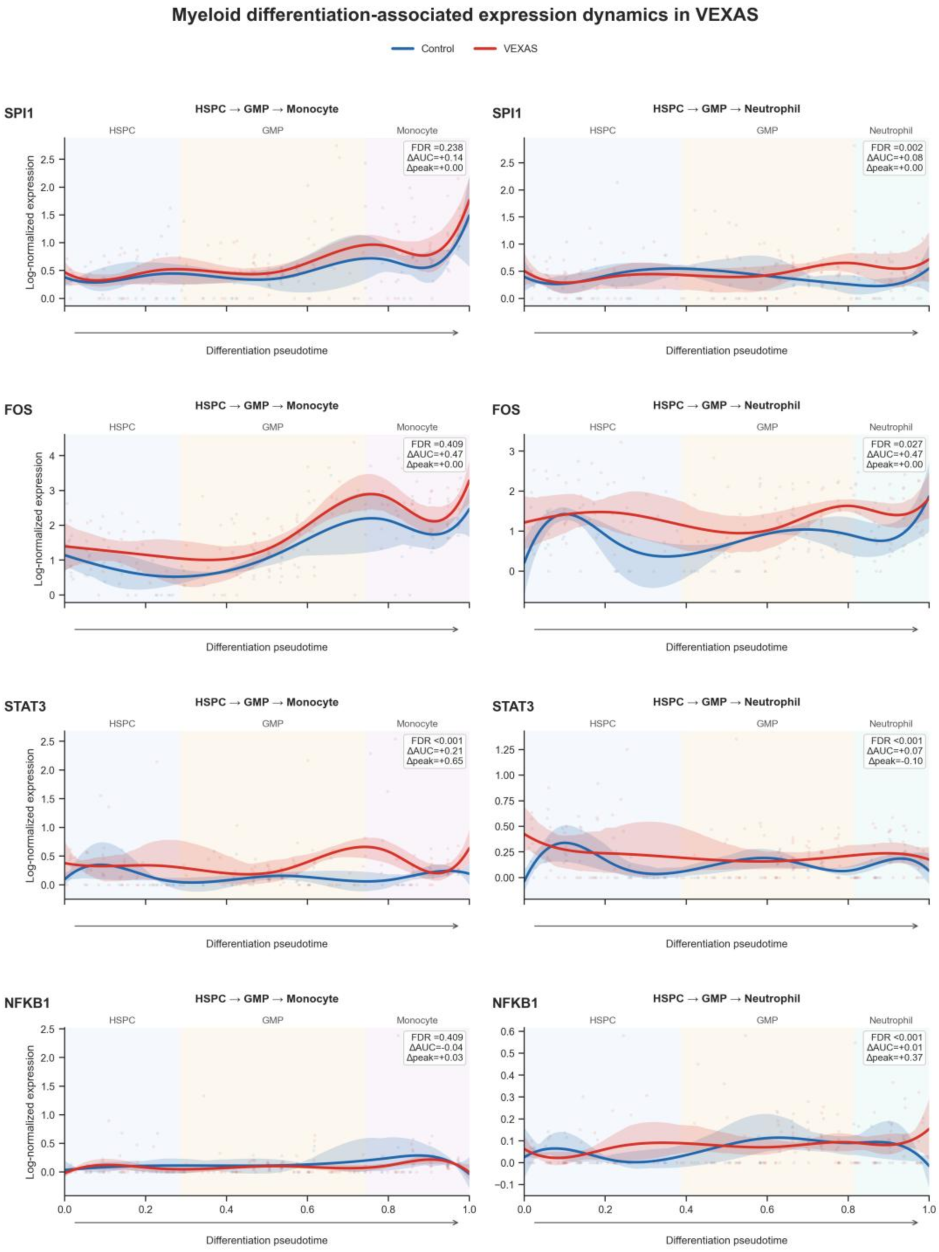

**Supplementary Figure S10: Pseudotime expression dynamics of representative cascade regulators.**

Smoothed log-normalized expression of SPI1, FOS, STAT3, and NFKB1 in control and VEXAS cells along HSPC-GMP-monocyte and HSPC-GMP-neutrophil trajectories. Shaded bands show uncertainty around the fitted curves, background shading marks cellular stages, and insets report branch-level FDR, difference in area under the curve, and difference in peak expression.

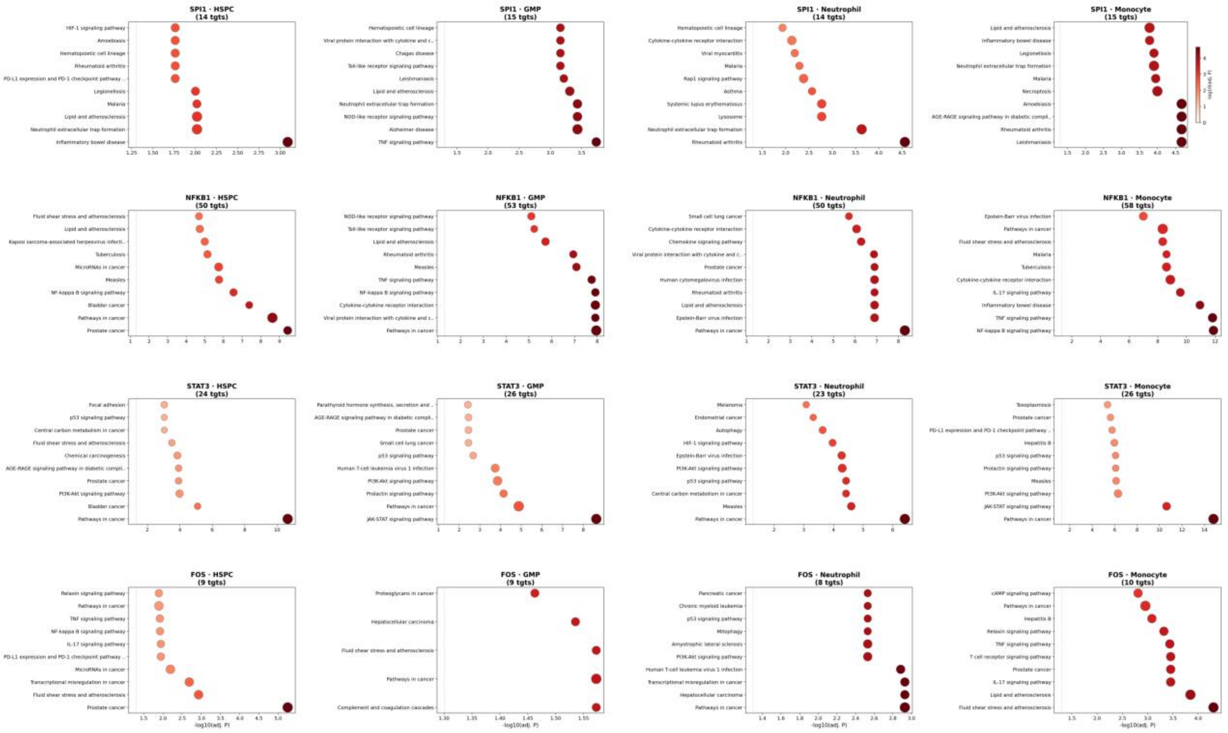

**Supplementary Figure S11: Cell type-specific functional enrichment of SPI1, NFKB1, STAT3, and FOS target sets.**

KEGG enrichment results for curated targets of the four prioritized transcription factors in HSPCs, GMPs, neutrophils, and monocytes. Point position and color denote -log10 adjusted P value, and panel titles report the number of targets evaluated in each transcription-factor/cell type combination.

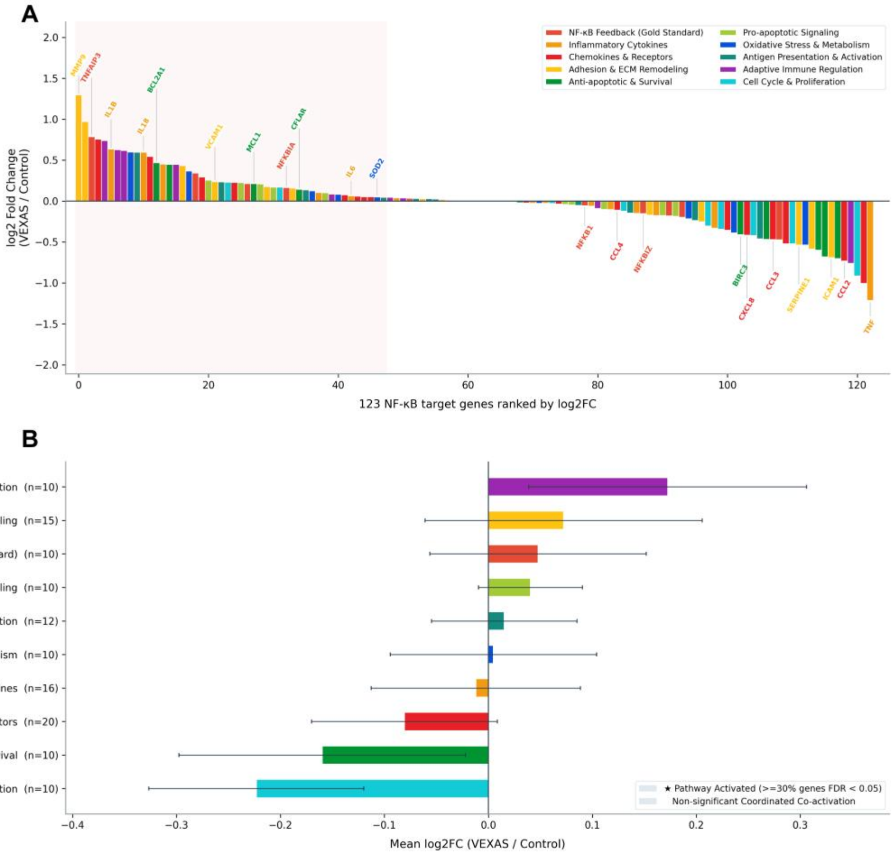

**Supplementary Figure S12: Expression landscape and functional organization of NFKB1 targets.**

**(A)** Log2 fold changes for 123 curated NFKB1 targets in VEXAS relative to control, ranked from increased to decreased expression and colored by functional category. Selected genes are labeled. **(B)** Mean log2 fold changes for functional target groups with dispersion intervals. Stars denote groups meeting the prespecified coordinated-activation criterion.

#### Supplementary Tables

**Supplementary Table 1: single-cell RNA-seq datasets used for IPF representation-learning and AML regulator-prioritization benchmarks.**

| GEO ID | Source | Condition | Sequencing Technology | Sample_num |
|---|---|---|---|---|
| GSE286182 | Lung | Control | 10X 3prime v3.1 | 30 |
| GSE286182 | Lung | IPF | 10X 3prime v3.1 | 24 |

| Sample Status | Sample ID | GEO ID | Cells per Sample | Total Samples |
|---|---|---|---|---|
| AML | AML1012 | GSM3587923 | 1136 | |
| AML | AML210A | GSM3587925 | 748 | |
| AML | AML314 | GSM3587927 | 162 | |
| AML | AML328 | GSM3587931 | 1094 | |
| AML | AML329 | GSM3587940 | 525 | |
| AML | AML371 | GSM3587946 | 756 | |
| AML | AML419A | GSM3587950 | 1189 | |
| AML | AML420B | GSM3587953 | 485 | 16 |
| AML | AML475 | GSM3587959 | 423 | |
| AML | AML556 | GSM3587963 | 2328 | |
| AML | AML707B | GSM3587969 | 1586 | |
| AML | AML722B | GSM3587980 | 79 | |
| AML | AML870 | GSM3587984 | 345 | |
| AML | AML916 | GSM3587988 | 933 | |
| AML | AML921A | GSM3587990 | 3813 | |
| AML | AML997 | GSM3587992 | 83 | |
| Healthy donor | BM1 | GSM3587996 | 108 | |
| Healthy donor | BM2 | GSM3587997 | 188 | |
| Healthy donor | BM3 | GSM3587998 | 643 | 6 |
| Healthy donor | BM4 | GSM3588000 | 3738 | |
| Healthy donor | BM5-34p | GSM3588002 | 1431 | |
| Healthy donor | BM5-34p38n | GSM3588003 | 1590 | |

Notes: GEO accession, tissue, disease status, sequencing platform, sample count, and cell count for GSE286182 and GSE116256.

**Supplementary Table 2: Independently curated AML reference genes for benchmarking ground truth gene list.**

| Gene name | Evidence | References (PMID) | Database Source |
|---|---|---|---|
| RUNX1 | High | 35957668,38104314 | DisGeNET & DepMap & Literature |
| CEBPA | High | 29956082,38104314 | DisGeNET & DepMap & Literature |
| SPI1 | High | 29956082 | DisGeNET & DepMap & Literature |
| CBFB | High | 27798625 | DisGeNET & DepMap & Literature |
| CDK6 | High | 33425766,25053825 | DisGeNET & DepMap & Literature |
| MYC | High | 29956082 | DisGeNET & DepMap & Literature |
| HOXA9 | Medium | 38104314 | DisGeNET & Literature |
| STAT3 | Medium | 29956082,37798266 | DisGeNET & Literature |
| GATA2 | Medium | 35957668 | DisGeNET & Literature |
| CCND2 | Medium | 27798625 | DisGeNET & Literature |
| S100A8 | Medium | 27798625 | DisGeNET & Literature |
| BCL2 | Medium | 33754282 | DisGeNET & Literature |
| JAK2 | Medium | 23970018 | DisGeNET & Literature |
| KMT2A | Medium | 26237430,10339604 | DisGeNET & Literature |
| EZH2 | Medium | 27798625 | DisGeNET & Literature |
| BCOR | Medium | 39936576 | DisGeNET & Literature |
| RUNX1T1 | Medium | 27798625 | DisGeNET & Literature |
| WT1 | Medium | 32171751 | DisGeNET & Literature |
| DDX41 | Medium | 32171751,39936576 | DisGeNET & Literature |
| MYH11 | Medium | 27798625,18206229 | DisGeNET & Literature |
| MECOM | Medium | 39936576 | DisGeNET & Literature |
| MYB | Medium | 38104314 | DepMap & Literature |
| LMO2 | Medium | 37573405 | DepMap & Literature |
| ZEB2 | Medium | 27756750 | DepMap & Literature |
| ERG | Medium | 19487285 | DisGeNET & Literature |
| ZEB1 | Low | 33960640,34550965 | Literature |
| MEIS1 | Low | 27335278 | Literature |
| ASXL1 | Low | 27335278 | Literature |
| ASXL2 | Low | 27335278 | Literature |
| BRD4 | Low | 41162275 | Literature |
| PBX3 | Low | 35957668 | Literature |
| MEF2C | Low | 35957668 | Literature |
| JUN | Low | 27840425 | Literature |

| | | | |
|---|---|---|---|
| TP53 | Low | 25822087 | DisGeNET (score > 0.9) |
| LYL1 | Low | 39464702 | DisGeNET (score > 0.9) |
| GFI1 | Low | 15457180 | DisGeNET (score > 0.9) |

Notes: The 36 AML reference genes, evidence level, supporting PMID(s), and contributing evidence source(s), including DepMap, DisGeNET, published studies, and expert review.

**Supplementary Table 3: Top 20 AML candidate regulators prioritized by CauNagi.**

| Rank | Gene | Score | High-confidence | Low-confidence | Irrelevant |
|---|---|---|---|---|---|
| 0 | STAT3 | 7.1059 | √ | | |
| 1 | GATA2 | 6.9995 | √ | | |
| 2 | STAT1 | 6.8973 | | | √ |
| 3 | SPI1 | 6.7432 | √ | | |
| 4 | RUNX1 | 6.7209 | √ | | |
| 5 | RBBP5 | 6.6248 | | | √ |
| 6 | TP53 | 6.5800 | | √ | |
| 7 | ZNF143 | 6.4351 | | | √ |
| 8 | GATA3 | 6.4124 | | | √ |
| 9 | ZEB1 | 6.2333 | | √ | |
| 10 | ZBTB7A | 5.9155 | | | √ |
| 11 | STAT4 | 5.7565 | | | √ |
| 12 | CHD7 | 5.7263 | | | √ |
| 13 | ATF3 | 5.7188 | | | √ |
| 14 | RXRA | 5.6086 | | | √ |
| 15 | JAK2 | 5.1016 | √ | | |
| 16 | CEBPA | 4.8675 | √ | | |
| 17 | EOMES | 4.7861 | | | √ |
| 18 | PRDM1 | 4.7751 | | | √ |
| 19 | ATF1 | 4.7692 | | | √ |

Notes: Top 20 candidate regulators ranked by CauNagi score. Genes were categorized as high-confidence, low-confidence, or irrelevant according to the AML reference annotation used for benchmarking. Check marks indicate category membership.

**Supplementary Table 4: Multi-source VEXAS single-cell RNA-seq cohorts and sample composition.**

| Dataset ID | Source | Disease Status | Sample count | Total Cells |
|---|---|---|---|---|
| GSE196052 | Bone Marrow | Healthy | 4 | 32247 |
| GSE196052 | Bone Marrow | VEXAS | 9 | 61280 |
| GSE216548 | PBMC | Healthy | 2 | 11260 |
| GSE216548 | PBMC | VEXAS | 2 | 12767 |
| GSE249131 | PBMC | Healthy | 5 | 54373 |
| GSE249131 | PBMC | VEXAS | 9 | 129759 |
| GSE272816 | Bone Marrow | VEXAS | 6 | 64061 |
| NCDATA | Bone Marrow | Healthy | 3 | 21487 |

Notes: Dataset accession, tissue source, disease status, number of donors, and available cell numbers for GSE196052, GSE216548, GSE249131, GSE272816, and the in-house healthy bone marrow cohort.

**Supplementary Table 5: Annotation and cascade metrics of 36 VEXAS cascade-core candidate regulators.**

| Gene | Vexas mechanism | Evidence level | CTS | HPS | LC |
|---|---|---|---|---|---|
| CEBPB | Myeloid Bias | Direct Validation | 1.2E-04 | 8.4E-01 | 9.8E-01 |
| CTCF | Other | Computational Prediction | 1.9E-04 | 8.2E-01 | 9.8E-01 |
| E2F1 | Other | Computational Prediction | 2.1E-03 | 6.5E-01 | 9.2E-01 |
| EBF1 | Lymphoid Depletion | Computational Prediction | 4.9E-04 | 7.2E-01 | 9.6E-01 |
| EGR1 | Other | Computational Prediction | 3.3E-03 | 6.8E-01 | 9.1E-01 |
| EP300 | Other | Computational Prediction | 6.8E-04 | 6.2E-01 | 9.6E-01 |
| FOS | UPR Stress | Indirect Support | 4.4E-04 | 7.8E-01 | 9.6E-01 |
| FOXA1 | Other | Computational Prediction | 1.5E-04 | 8.0E-01 | 9.8E-01 |
| FOXA2 | Other | Computational Prediction | 2.0E-03 | 7.3E-01 | 9.3E-01 |
| GATA1 | Erythroid Defect | Computational Prediction | 6.0E-04 | 6.5E-01 | 9.6E-01 |
| GATA2 | Other | Computational Prediction | 4.9E-05 | 8.0E-01 | 9.9E-01 |
| GATA3 | Other | Computational Prediction | 2.4E-04 | 6.5E-01 | 9.7E-01 |
| HEY1 | Other | Computational Prediction | 1.7E-03 | 6.8E-01 | 9.3E-01 |
| IRF4 | Other | Computational Prediction | 2.6E-03 | 6.2E-01 | 9.1E-01 |
| JUN | UPR Stress | Indirect Support | 1.0E-03 | 7.7E-01 | 9.5E-01 |
| JUND | UPR Stress | Indirect Support | 1.3E-04 | 8.3E-01 | 9.8E-01 |
| MAFF | Other | Computational Prediction | 1.1E-03 | 7.6E-01 | 9.4E-01 |
| MAFK | Other | Computational Prediction | 3.4E-04 | 8.1E-01 | 9.7E-01 |
| MAX | Other | Computational Prediction | 2.4E-04 | 7.2E-01 | 9.7E-01 |
| MEF2A | Other | Computational Prediction | 1.5E-03 | 6.7E-01 | 9.4E-01 |
| MYC | Other | Computational Prediction | 1.9E-03 | 6.9E-01 | 9.3E-01 |
| NFKB1 | NFkB-Signaling | Direct Validation | 4.1E-04 | 7.0E-01 | 9.7E-01 |
| PAX5 | Lymphoid Depletion | Direct Validation | 1.9E-03 | 7.0E-01 | 9.3E-01 |
| POL2 | Other | Computational Prediction | 7.7E-04 | 6.1E-01 | 9.5E-01 |
| POLR2A | Other | Computational Prediction | 1.3E-04 | 6.2E-01 | 9.8E-01 |
| RAD21 | Other | Computational Prediction | 8.2E-05 | 8.0E-01 | 9.8E-01 |
| SMC3 | Other | Computational Prediction | 2.6E-04 | 7.4E-01 | 9.7E-01 |
| SPI1 | Myeloid Bias | Direct Validation | 2.2E-04 | 8.2E-01 | 9.8E-01 |
| STAT3 | JAK-STAT | Therapeutic Evidence | 1.9E-04 | 7.9E-01 | 9.8E-01 |
| TAF1 | Other | Computational Prediction | 2.9E-04 | 7.8E-01 | 9.7E-01 |
| TAL1 | Other | Computational Prediction | 7.2E-04 | 6.4E-01 | 9.6E-01 |
| TBP | Other | Computational Prediction | 3.2E-04 | 7.6E-01 | 9.7E-01 |
| TCF12 | Lymphoid Depletion | Computational Prediction | 2.4E-02 | 8.2E-01 | 7.5E-01 |
| TCF4 | Lymphoid Depletion | Computational Prediction | 4.6E-03 | 6.9E-01 | 8.9E-01 |
| USF1 | Other | Computational Prediction | 8.2E-04 | 7.9E-01 | 9.5E-01 |
| YY1 | Other | Computational Prediction | 7.7E-04 | 7.9E-01 | 9.5E-01 |

Notes: Gene-level annotation of the 36 cascade-core candidate regulators identified by CauNagi, including their putative VEXAS-associated mechanisms, evidence levels, cascade transition specificity (CTS), hierarchical propagation score (HPS), and layer consistency (LC). Evidence levels summarize available experimental, therapeutic, indirect, or computational support for each candidate.

**Supplementary Table 6: Human bulk RNA-seq samples used for external validation.**

| Dataset ID | Source | Disease Status | Sample count | Sequencing Platform |
|---|---|---|---|---|
| GSE272578 | HSPCs, 24 h post-editing | Healthy | 2 | Illumina NovaSeq 6000 |
| GSE272578 | HSPCs, 24 h post-editing | VEXAS | 3 | Illumina NovaSeq 6000 |
| GSE272578 | HSPCs, 7 d post-editing | Healthy | 3 | Illumina NovaSeq 6000 |
| GSE272578 | HSPCs, 7 d post-editing | VEXAS | 3 | Illumina NovaSeq 6000 |
| GSE272578 | Monocytes | Healthy | 3 | Illumina NovaSeq 6000 |
| GSE272578 | Monocytes | VEXAS | 5 | Illumina NovaSeq 6000 |

Notes: Sample source, experimental state, collection time, sample count, and sequencing platform for UBA1-edited HSPCs and VEXAS/control monocytes in GSE272578.
